# MOSTWAS: Multi-Omic Strategies for Transcriptome-Wide Association Studies

**DOI:** 10.1101/2020.04.17.047225

**Authors:** Arjun Bhattacharya, Yun Li, Michael I. Love

**Affiliations:** Department of Biostatistics, University of North Carolina at Chapel Hill, Chapel Hill, NC, USA; Department of Genetics, University of North Carolina at Chapel Hill, Chapel Hill, NC, USA; Department of Computer Science, University of North Carolina at Chapel Hill, Chapel Hill, NC, USA

**Keywords:** transcriptome-wide association study, genome-wide association study, expression quantitative trait loci analysis, multi-omics, mediation analysis

## Abstract

Traditional predictive models for transcriptome-wide association studies (TWAS) consider only single nucleotide polymorphisms (SNPs) local to genes of interest and perform parameter shrinkage with a regularization process. These approaches ignore the effect of distal-SNPs or other molecular effects underlying the SNP-gene association. Here, we outline multi-omics strategies for transcriptome imputation from germline genetics to allow more powerful testing of gene-trait associations by prioritizing distal-SNPs to the gene of interest. In one extension, we identify mediating biomarkers (CpG sites, microRNAs, and transcription factors) highly associated with gene expression and train predictive models for these mediators using their local SNPs. Imputed values for mediators are then incorporated into the final predictive model of gene expression, along with local SNPs. In the second extension, we assess distal-eQTLs (SNPs associated with genes not in a local window around it) for their mediation effect through mediating biomarkers local to these distal-eSNPs. Distal-eSNPs with large indirect mediation effects are then included in the transcriptomic prediction model with the local SNPs around the gene of interest. Using simulations and real data from ROS/MAP brain tissue and TCGA breast tumors, we show considerable gains of percent variance explained (1-2% additive increase) of gene expression and TWAS power to detect gene-trait associations. This integrative approach to transcriptome-wide imputation and association studies aids in identifying the complex interactions underlying genetic regulation within a tissue and important risk genes for various traits and disorders.

**AUTHOR SUMMARY:** Transcriptome-wide association studies (TWAS) are a powerful strategy to study gene-trait associations by integrating genome-wide association studies (GWAS) with gene expression datasets. TWAS increases study power and interpretability by mapping genetic variants to genes. However, traditional TWAS consider only variants that are close to a gene and thus ignores important variants far away from the gene that may be involved in complex regulatory mechanisms. Here, we present MOSTWAS (Multi-Omic Strategies for TWAS), a suite of tools that extends the TWAS framework to include these distal variants. MOSTWAS leverages multi-omic data of regulatory biomarkers (transcription factors, microRNAs, epigenetics) and borrows from techniques in mediation analysis to prioritize distal variants that are around these regulatory biomarkers. Using simulations and real public data from brain tissue and breast tumors, we show that MOSTWAS improves upon traditional TWAS in both predictive performance and power to detect gene-trait associations. MOSTWAS also aids in identifying possible mechanisms for gene regulation using a novel added-last test that assesses the added information gained from the distal variants beyond the local association. In conclusion, our method aids in detecting important risk genes for traits and disorders and the possible complex interactions underlying genetic regulation within a tissue.

## INTRODUCTION

Genomic methods that borrow information from multiple data sources, or “omics” assays, offer advantages in interpretability, statistical efficiency, and opportunities to understand causal molecular pathways in disease regulation [1,2]. Transcriptome-wide associations studies (TWAS) aggregate genetic information into functionally relevant testing units that map to genes and their expression in a trait-relevant tissue. This gene-based approach combines the effects of many regulatory variants into a single testing unit that can increase study power and aid in interpretability of trait-associated genomic loci [3,4]. However, traditional TWAS methods, like PrediXcan [3] and FUSION [4], focus on local genetic regulation of transcription. These methods ignore significant portions of heritable expression that can be attributed to distal genetic variants that may indicate complex mechanisms contributing to gene regulation.

Recent work in transcriptional regulation has estimated that distal genetic traits account for up to 70% of the variance in gene expression [5,6]. These results accord with Boyle *et al*’s omnigenic model, proposing that regulatory networks are so interconnected that a majority of genetic variants in the genome, local or distal, have indirect effects on the expression level of any particular gene [6,7]. In fact, work by Sinnott-Armstrong *et al* showed huge enrichment of significant genetic signal near genes involved in the relevant pathways for biologically simple traits, even for phenotypes largely thought to be simpler than complex diseases [8]. Together, these observations suggest that the majority of phenotype variance, even for traits commonly believed to be simpler than complex diseases like cancer, is not driven by variants in core genes, but rather from thousands of variants spreading across most of the genome.

Many groups have leveraged the omnigenic model to identify distal expression quantitative loci (eQTLs) by testing the effect of a distal-eSNP on an gene mediated through a set of genes local to the distal-eSNP, where the SNP and gene are more than 0.5 Megabases (Mb) away. These studies draw the conclusion that many distal-eQTLs are often eQTLs for one of more of their local genes [9–15]. It has been shown previously that distal-eQTLs found in regulatory hotspots are often cell-type specific [9,13,16] and hence carry biologically relevant signal when studying bulk tissue with heterogeneous cell-types (e.g. cancerous tumors or the brain). More recently, the concepts of distal-eQTLs residing in or near regulatory elements have been integrated with multi-omics data and biological priors to reconstruct molecular networks and hypothesize cell-regulatory mechanisms [17].

Variant-mapping methods have also shown the utility of integrating molecular data beyond transcriptomics. Deep learning methods have been employed to link GWAS-identified variants to nearby regulatory mechanisms to generate functional hypotheses for SNP-trait associations [18–20]. These ideas have been extended to TWAS: the EpiXcan method demonstrates that incorporating epigenetic information into transcriptomic prediction models generally improves predictive performance and power in detecting gene-trait associations in local-only TWAS [21]. Wheeler *et al* have leveraged TWAS imputation to show that *trans*-acting genes are often found in transcriptional regulation pathways and are likely to be associated with complex traits [22]. Thus, it is imperative to prioritize distal variants that are *trans*-acting to fully capture heritable gene expression that is associated with complex diseases in TWAS.

To this end, we developed two extensions to TWAS, borrowing information from other omics assays to enrich or prioritize mediator relationships of eQTLs in expression models. Using simulations and data from Religious Orders Study and the Rush Memory and Aging Project (ROS/MAP) [23] and The Cancer Genome Atlas (TCGA) [24], we show considerable improvements in transcriptomic prediction and power to detect gene-trait associations. These **M**ulti-**O**mic **S**trategies for **T**ranscriptome-**W**ide **A**ssociation **S**tudies are curated in the R package MOSTWAS, available freely at https://bhattacharya-a-bt.github.io/MOSTWAS.

## RESULTS

### Overview of MOSTWAS

MOSTWAS incorporates two methods to include distal-eQTLs in transcriptomic prediction: mediator-enriched TWAS (MeTWAS) and distal-eQTL prioritization via mediation analysis (DePMA). Here, we refer to an eQTL as a SNP with an association with the expression of a gene, and a distal-eQTL is more than 0.5 Mb away from the eGene. As large proportions of total heritable gene expression are explained by distal-eQTLs local to regulatory hotspots [6,11,13,14], we used data-driven approaches to either identify mediating regulatory biomarkers (MeTWAS) or distal-eQTLs mediated by local biomarkers (DePMA) to increase predictive power for gene expression and power to detect gene-trait associations. These methods are described in **Methods** with an algorithmic summary in **Supplemental Figure S1**.

**Figure 1** provides an example of the biological mechanisms MOSTWAS attempts to leverage in its predictive models for a gene *G* of interest: here, without loss of generality of the regulatory mechanism, assume a SNP within a regulatory element affects the transcription of gene *X* that codes for a transcription factor. Transcription factor X then binds to a distal regulatory region and affects the transcription of gene *G*. Methodologically,

**Figure 1:**
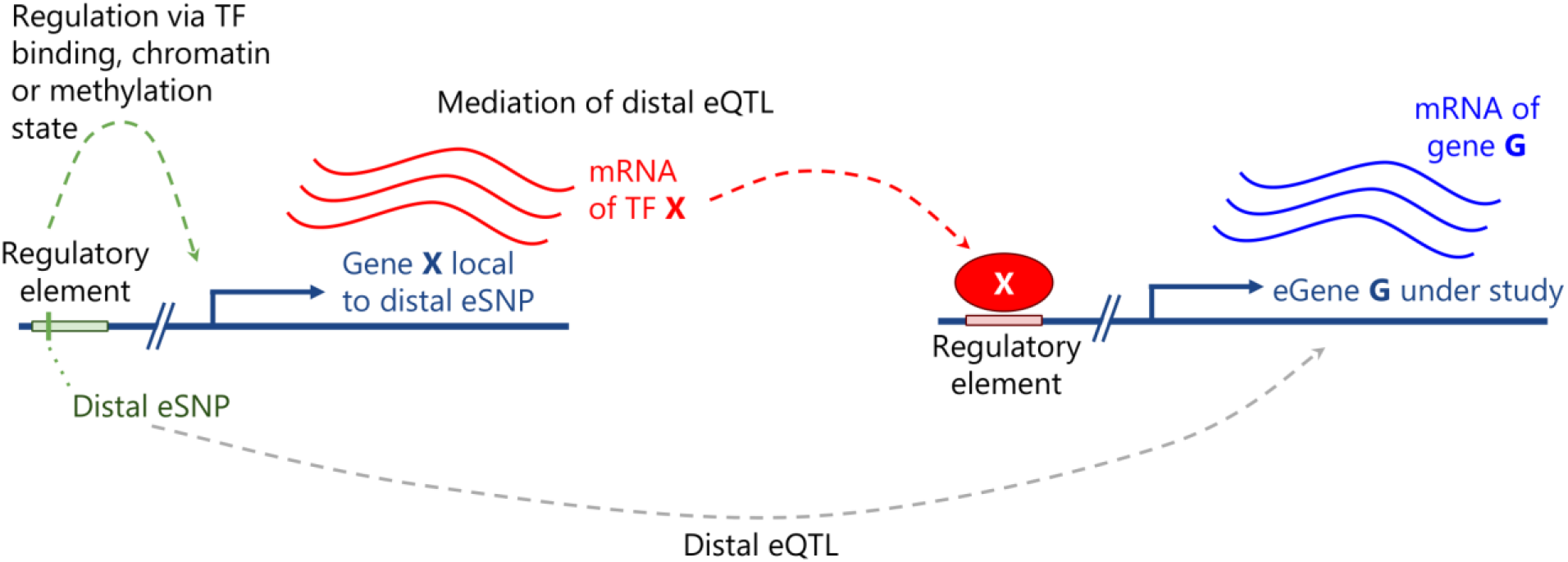
Example of a biological mechanism MOSTWAS leverages in its predictive models. Here, assume a SNP (in green) within a regulatory element affects the transcription of gene *X* that codes for a transcription factor. Transcription factor *X* then binds to a distal regulatory region and affects the transcription of gene *G*. The association between the expression of gene *X* and gene *G* is leveraged in the first step of MeTWAS. A distal-eQTL association is also conferred between this distal-SNP and the eGene *G*, which is leveraged in the DePMA training process.

- MeTWAS first detects the association between the expression of gene *X* and expression of gene *G*. It proceeds upstream in the regulatory pathway to the genetic locus around gene *X* and builds a predictive model for the expression of gene *X*. Imputed expression of gene *X* (imputed via cross-validation) is then included as a fixed effect in the predictive model of gene *G*, along with the genetic variants local to gene *G*. This model is fit using a two-stage regression model [25], first fitting the imputed mediators using least squares regression and then the local genotypes using elastic net regression [26] or linear mixed modeling [27].
- DePMA first detects the distal-eQTL association between the distal SNP and expression of gene *G*. It then proceeds downstream in the regulatory pathway from the distal SNP to identify whether there is a strong association between the SNP and the expression of the local gene *X*. Using mediation analysis, if the indirect effect of the SNP on gene *G* mediated through gene *X* is significantly large, the SNP is included in the final predictive model for the expression of gene *G*, fit using elastic net regression [26] or linear mixed modeling [27].

MeTWAS and DePMA can consider any set of regulatory elements as potential mediators (e.g. transcription factors, microRNAs, CpG methylation sites, chromatin-binding factors, etc.).

If individual genotype data is available in an external GWAS panel, a MeTWAS or DePMA model may be used to impute tissue-specific expression. If only summary statistics are available in the GWAS panel, the Imp-G weighted burden testing framework [28] as implemented in FUSION [4] can be applied. We further implement a permutation test to assess whether the overall gene-trait association is significant, conditional on the GWAS effect sizes [4] and a novel distal-SNPs added-last test that assesses the added information from distal-SNPs given the association from the local SNPs (**Methods**).

### Simulation analysis

We first conducted simulations to assess the power to predict gene expression and power to detect gene-trait associations under various settings for phenotype heritability, local heritability of expression, distal heritability of expression, and proportion of causal local and distal SNPs for MeTWAS and DePMA (full simulation details in **Methods**). Using genetic data from TCGA-BRCA as a reference, we used SNPs local to the gene *ESR1* (Chromosome 6) to generate local eQTLs and SNPs local to *FOXA1* (Chromosome 14) to generate distal-eQTLs for a 400-sample eQTL reference panel and 1,500-sample GWAS imputation panel. We considered two scenarios for each set of simulation parameters: (1) an ideal case where the leveraged associated between the distal-SNP and gene of interest exists in both the reference and imputation panel, and (2) a “null” case where the leveraged association between the distal-SNP and the gene of interest exists in the reference panel but does not contribute to phenotype heritability in the imputation panel. Though the choice of these loci was arbitrary for constructing the simulation, there is evidence that *ESR1* and *FOXA1* are highly co-expression in breast tumors, and local-eQTLs of *FOXA1* have been shown to be distal-eQTLs of *ESR1* [29].

In these simulation studies, we found that MOSTWAS methods performed well in prediction across different causal proportions and local and distal mRNA expression heritabilities and generally outperform local-only modelling. Furthermore, across all simulation settings, we observed that MOSTWAS showed greater or nearly equal power to detect gene-trait associations compared to local-only models. We found that, under the setting that distal-eQTLs contributes to trait heritability, the best MOSTWAS model had greater power to detect gene-trait associations than the local-only models, with the advantage in power over local-only models increasing with increased distal expression heritability (**Figure 2A**). Similarly, we found that as the proportion of total expression heritability attributed to distal variation increased, the positive difference in predictive performance between the best MOSTWAS model and the local-only model increased (**Supplemental Figure S2**). Under the “null” case that distal variation influences expression only in the reference panel, as expected, we observed that local-only and MOSTWAS models perform similarly. Only at low causal proportions (causal proportion of 0.01) and low trait heritability (trait heritability of 0.2), did local-only models have a modest advantage in TWAS power over MOSTWAS models (**Figure 2B** and **Supplemental Data**). This difference was reduced at larger causal proportions and trait heritabilities (**Figure 2B**). Using these same simulation parameters, we also simulated the false positive rate (FPR), defined as the proportion of positive associations at *P* < 0.05 under the null, where the phenotype trait in the GWAS panel was permuted 1,000 times across 20 sets of simulations. We found that the FPR was generally around 0.05 for all methods (**Supplemental Figure S3**).

**Figure 2:**
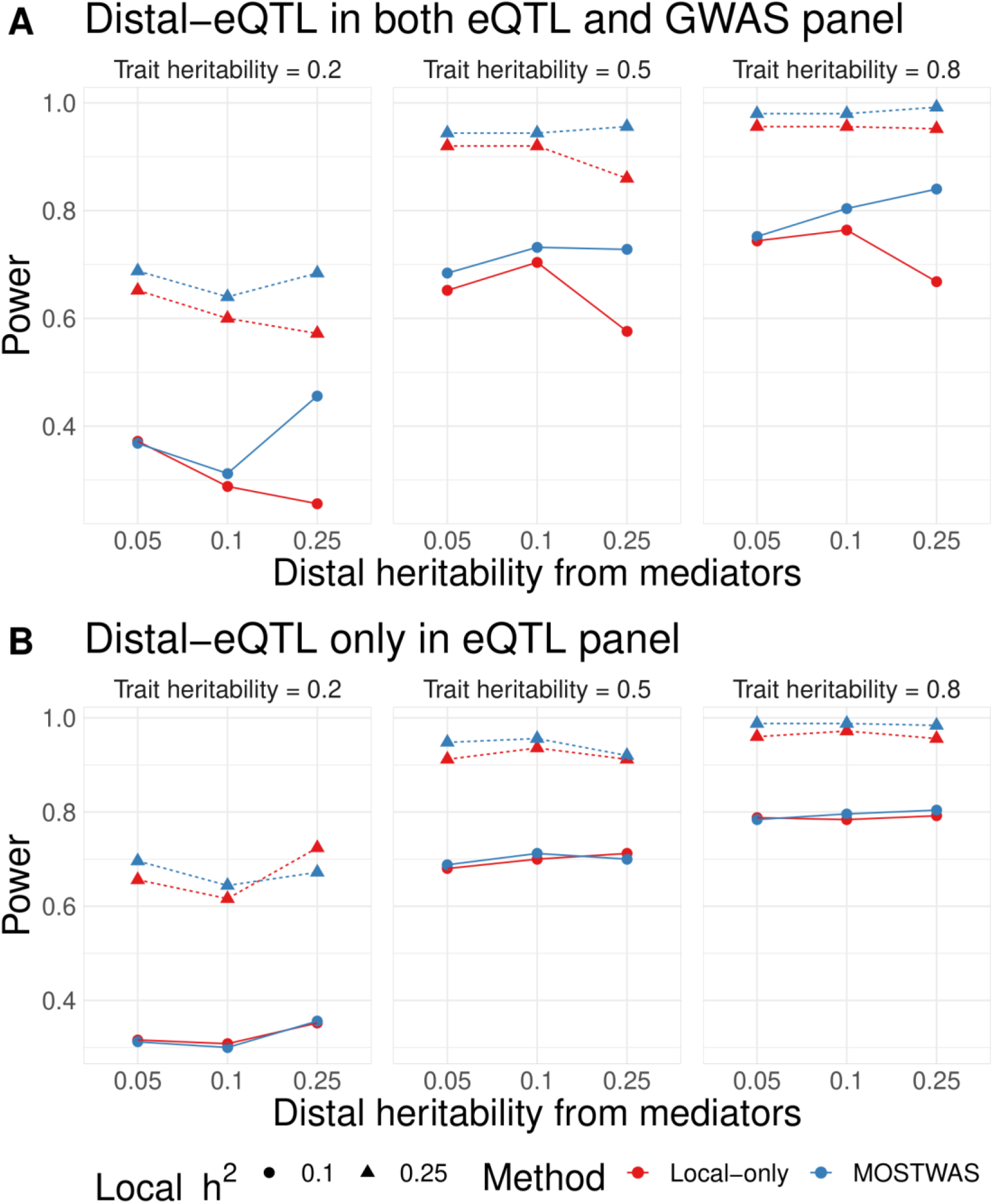
Comparison of TWAS power via simulations using MOSTWAS and local-only models. **(A)** Proportion of gene-trait associations at *P* < 2.5 × 10^−6^ using local-only (red) and the most predictive MOSTWAS (blue) models across various local and distal expression heritabilities, trait heritability, and causal proportions. **(B)** Proportion of significant gene-trait associations across the same simulation parameters with no distal effect on the trait in the simulated external GWAS panel.

The power of the distal-SNPs added-last test increased significantly as both the sample sizes of the eQTL reference panel and the GWAS imputation panel increased (**Supplemental Figure S4**). At a sample size of 10,000 in the GWAS panel with summary statistics (a suitably large GWAS) and a sample size greater than 200 in the eQTL panel, MOSTWAS obtained over 65% power to detect significant distal significant associations (**Supplemental Figure S4**). Overall, these results demonstrated the advantages of MOSTWAS methods for modeling the complex genetic architecture of transcriptomes, especially when distal variation has a large effect on the heritability of both the gene and trait of interest. Full simulation results are provided in **Supplemental Data**, accessible at https://zenodo.org/record/3755919 [30]. The MOSTWAS package also contains functions for replicating this simulation framework.

### Real data applications in brain tissue

We applied MOSTWAS to multi-omic data derived from samples of prefrontal cortex, a tissue that has been used previously in studying neuropsychiatric traits and disorders with TWAS [44,45]. There is ample evidence from studies of brain tissue, especially the prefrontal cortex, that non-coding variants may regulate distal genes [44,46,47]; in fact, an eQTL analysis by Sng *et al* found that approximately 20-40% of detected eQTLs in the frontal cortex can be considered *trans*-acting [48]. Thus, the prefrontal cortex in the context of neuropsychiatric disorders provides a prime example to assess MOSTWAS.

Using ROS/MAP data on germline SNPs, tumor mRNA expression, CpG DNA methylation, and miRNA expression (*N* = 370), we trained MeTWAS, DePMA, and traditional local-only predictive models for the tumor expression of all genes with significant non-zero heritability. Estimates of gene expression heritability were considerably larger when we considered distal variation with MOSTWAS (**Supplemental Table S1**). We also found that MeTWAS and DePMA performed better in cross-validation *R*^2^ than local-only models (**Figures 3A-C**). Mean predictive *R*^2^ for local-only models was 0.029 (25% to 75% inter-quartile interval (0.0,0.015)), for MeTWAS models was 0.079 (0.019, 0.082), and for DePMA models was 0.045 (0.013, 0.037).

**Figure 3:**
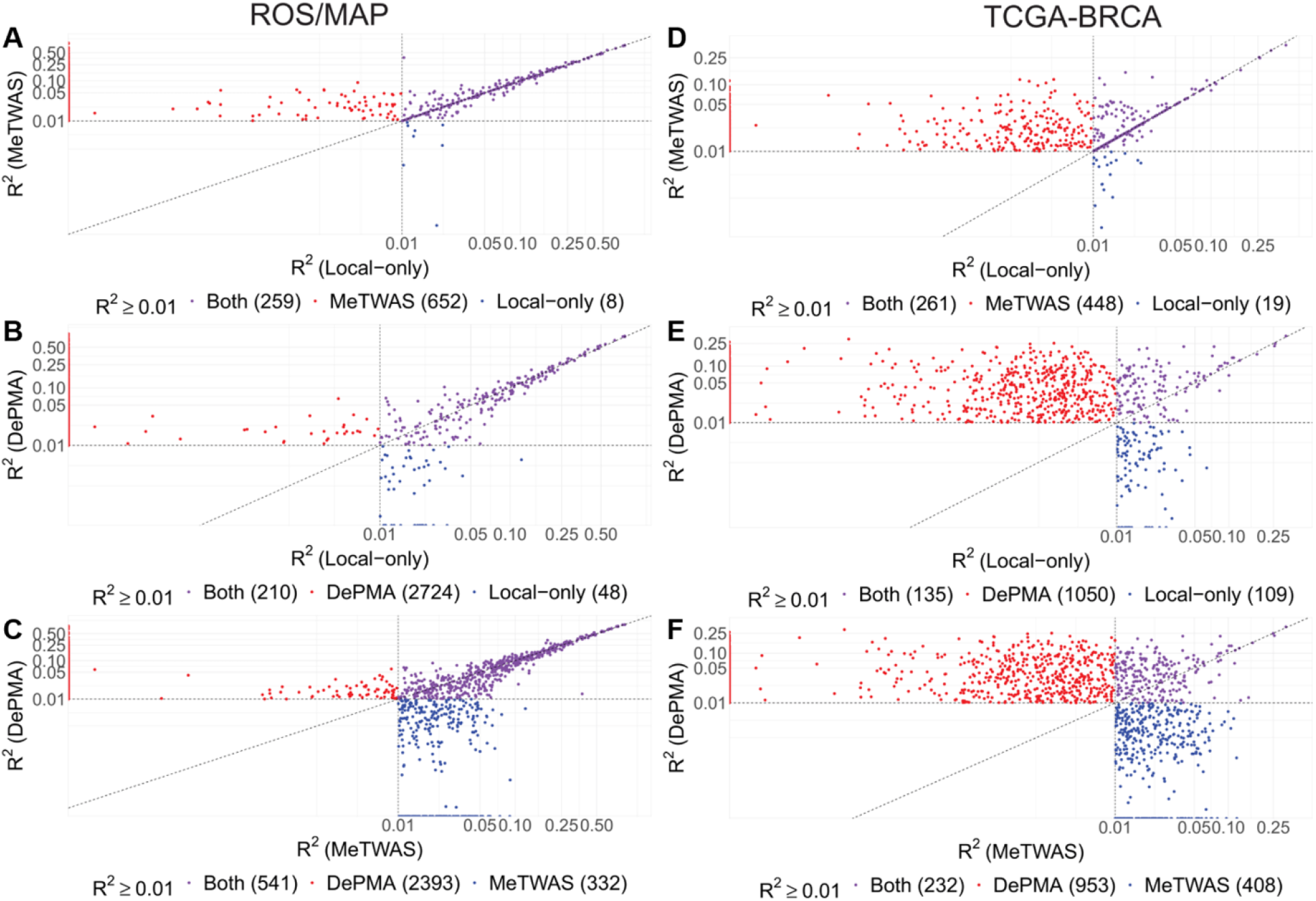
Predictive adjusted R^2^ from cross-validation across local-only, MeTWAS, and DePMA models. If a given gene does not have *h*^2^ > 0 with *P* < 0.05, we set the predictive adjusted *R*^2^ to 0 here for comparison. The top row compares local-only and MeTWAS, middle row compares local-only and DePMA, and the bottom row compares MeTWAS and DePMA. The left column has performance in ROS/MAP, while the right column has performance in TCGA-BRCA. All axes indicate the CV adjusted *R*^2^ for different models.

We used 87 samples in ROS/MAP with genotype and mRNA expression data that were not used in model training to test portability of MOSTWAS models in independent cohorts. As shown in **Figure 4A** and **Supplemental Figure S5**, DePMA models obtained the highest predictive adjusted *R*^2^ in the external cohort (0.042 (0.009, 0.057)), with MeTWAS (0.040 (0.010, 0.054)) also outperforming local-only models (0.031 (0.007, 0.039)). Overall, among genes with cross-validation adjusted *R*^2^ ≥ 0.01, 187 out of 267 genes achieved external predictive *R*^2^ ≥ 0.01 using local-only models, 683 out of 911 using MeTWAS, and 2,135 out of 2,934 using DePMA (**Figure 3A-C**).

**Figure 4:**
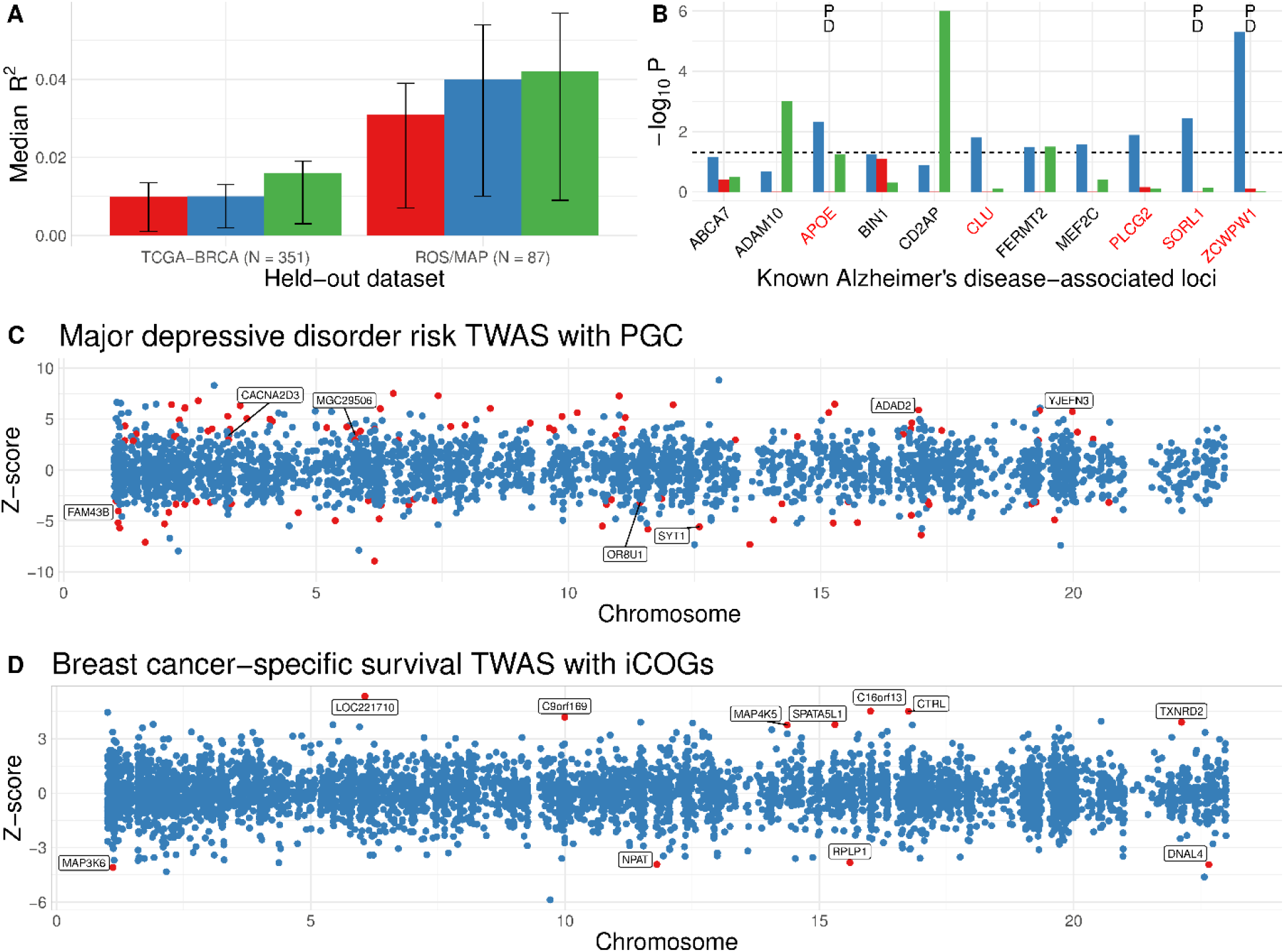
External validation of MOSTWAS and gene-trait associations using MOSTWAS models. **(A)** Median predictive adjusted $R^2$ in held-out cohorts from TCGA-BRCA and ROS/MAP in local-only, MeTWAS, and DePMA models that have in-sample significant heritability. The interval shows the 25% and 75% quantiles for external cohort predictive *R*^2^. **(B)** Associations with 11 known Alzheimer’s risk loci, as identified in literature, using MOSTWAS, local-only, and TIGAR Dirichlet process regression (DPR). Loci are labeled with P if the permutation test achieves FDR-adjusted *P* < 0.05 and D if the added-last test achieves FDR-adjusted *P* < 0.05. **(C)** TWAS associations for major depressive disorder risk using GWAS summary statistics from PGC. Loci are colored red if the overall association achieves FDR-adjusted *P* < 0.05 and the permutation test also achieves FDR-adjusted *P* < 0.05. We label the 7 loci that were independently validated with UK Biobank GWAX summary statistics at FDR-adjusted *P* < 0.05 for both the overall association test and permutation test. **(D)** TWAS associations for breast cancer-specific survival using GWAS summary statistics from iCOGs. Loci are colored red and labeled if the overall association achieves FDR-adjusted *P* < 0.05.

We next conducted association tests for known Alzheimer’s disease risk loci using local-only and the best MOSTWAS model (selected by comparing MeTWAS and DePMA cross-validation *R*^2^) trained in ROS/MAP and summary-level GWAS data from the International Genomics of Alzheimer’s Project (IGAP) [49]. From literature, we identified 14 known common and rare loci of late-onset Alzheimer’s disease that have been mapped to genes [49–52], 11 of which had MOSTWAS models with cross-validation *R*^2^ ≥ 0.01. Five of these 11 loci (*APOE*, *CLU*, *PLCG2*, *SORL1*, *ZCWPW1*) showed significant association at FDR-adjusted *P* < 0.05 (**Supplemental Table S2**). We also compared these 11 associations to those identified by local-only models (PrediXcan [3] and TIGAR [53]), with raw *P*-values of association shown in **Figure 4B**. MOSTWAS showed stronger associations at 8 of these loci than both local-only and DPR models. We followed up on the 5 significantly associated loci using the permutation and added-last tests (**Methods** and **Supplemental Methods**). Three of these loci (*APOE*, *SORL1*, *ZCWPW1*) showed significant associations, conditional on variants with large GWAS effect sizes (permutation test significant at FDR-adjusted *P* < 0.05). These three loci also showed significant associations with distal variants, above and beyond the association with local variants, at FDR-adjusted *P* < 0.05 (**Supplemental Table S2**).

We then conducted a transcriptome-wide association study for risk of major depressive disorder (MDD) using summary statistics from the Psychiatric Genomics Consortium (PGC) genome-wide meta-analysis that excluded data from the UK Biobank and 23andMe [54]. QQ-plots for TWAS *Z*-statistics and *P*-values are provided in **Supplemental Figure S7** and **Supplemental Figure S8** for both local-only and MOSTWAS models, showing earlier departure from the null using local-only models compared to MOSTWAS. Overall, using all heritable genes with cross-validation *R*^2^ with the best MOSTWAS model in ROS/MAP, we identified 102 MDD risk-associated loci with FDR-adjusted *P* < 0.05 that persisted when subjected to permutation testing at FDR-adjusted *P* < 0.05 (colored red in **Figure 4C**). We downloaded genome-wide association study by proxy (GWAX) summary statistics from the UK Biobank [55] for replication analysis of loci identified using PGC summary statistics. We found 7 of these 102 loci (labeled in **Figure 4C** and listed in **Supplemental Table S3**) also showed an association in UK Biobank GWAS that was in the same direction as in PGC. In comparison, using local-only models, we identified 11 genes with significant association with MDD risk at FDR-adjusted *P* < 0.05 that persisted after permutation testing; none of these loci showed significant associations in the UK Biobank GWAX in the same direction as in PGC. These replication rates between MOSTWAS and local-only models were similar (accounting for the total number of associations), highlighting that the inclusion of distal variation does not hinder the replicability of MOSTWAS associations in comparison to local-only models [55,56]. Local-only results are provided in **Supplemental Data**. It is important to note here that the UK Biobank dataset is not a GWAS dataset as it defined a case of MDD as any subject who has the disorder or a first-degree relative with MDD. Hence, the study forfeits study power to detect gene-trait associations for MDD [55,56]. Nonetheless, we believe that strong prediction in independent cohorts and TWAS results across two independent cohorts provided an example of the robustness of MOSTWAS models.

In summary, we observed that MOSTWAS models generally had higher predictive *R*^2^ than local-only models both in training and independent cohorts. We also found that MOSTWAS recapitulated 5 known Alzheimer’s risk loci that were not detected by local-only modeling (both PrediXcan [3] and TIGAR [53]), 3 of which had significant distal associations above and beyond the information in local variants using our added-last test. We also illustrated that some MDD-risk-associated loci detected by MOSTWAS in a GWAS cohort were replicable in an independent GWAX cohort [54,55].

### Real data applications in breast cancer tumors

We applied MOSTWAS using breast tumor multi-omics and disease outcomes, motivated by recent GWAS and TWAS for breast cancer-specific survival [31–35]. Previous breast tumor eQTL studies have revealed several significant distal-eQTLs in trait-associated loci, many of which are in regulatory or epigenetic hotspots [35,36], motivating our application of MOSTWAS in breast tumor expression modeling.

Using TCGA-BRCA [24] datasets for germline SNPs, tumor mRNA expression, CpG DNA methylation, and miRNA expression (*N* = 563), we trained MeTWAS, DePMA, and traditional local-only predictive models for the mRNA expression of all genes with significant non-zero germline heritability at *P* < 0.05. Estimates of heritability for genes were 2-4 times larger when we considered distal variation using MOSTWAS methods (**Supplemental Table S1**). We also found that MeTWAS and DePMA performed better in cross-validation *R*^2^, with larger numbers of models at *R*^2^ ≥ 0.01 and significant germline heritability using MOSTWAS models than local-only models (**Figures 3D-F**). Mean predictive *R*^2^ for local-only models was 0.011 (25% to 75% inter-quartile interval (0.0,0.013)), for MeTWAS models was 0.028 (0.013, 0.032), and for DePMA models was 0.051 (0.019, 0.068).

In addition to cross-validation, we used 351 samples in TCGA-BRCA with only genotype and mRNA expression data, which were not used in model training, to test the portability of MOSTWAS models in independent external cohorts. As shown in **Figure 4A** and **Supplemental Figure S5**, DePMA models obtained the highest predictive adjusted *R*^2^ in the external cohort (mean 0.016, 25% to 75% inter-quartile interval (0.003,0.018)), with MeTWAS models (0.011, (0.002,0.014)) performing on par with local-only models (0.010, (0.001, 0.015)), considering only genes that showed significant heritability and cross-validation adjusted *R*^2^ ≥ 0.01 using a given method. Overall, among genes with cross-validation adjusted R^2^ ≥ 0.01, 37 out of 280 achieved external predictive *R*^2^ ≥ 0.01 using local-only models, 89 out of 709 using MeTWAS, and 787 out of 1,185 using DePMA (**Figure 3D-F**).

Lastly, we conducted association studies for breast cancer-specific survival using local-only and the MOSTWAS model with largest *R*^2^ trained in TCGA-BRCA and summary-level GWAS data from iCOGs [34]. Here, we constructed the weight burden test, as described above and in Pasaniuc *et al* and Gusev *et al* [4,28]. We prioritized genes with Benjamini-Hochberg (BH) [37] adjusted *P* < 0.05 for permutation testing. Of the 122 genes that had cross-validation *R*^2^ ≥ 0.01 in TCGA-BRCA using both local-only and MOSTWAS models, we found 2 survival associations with the same loci at BH FDR-adjusted *P* < 0.05, with the strength of association marginally larger with the MOSTWAS model in each case (**Supplemental Figure S6**). Furthermore, 115 of these loci showed larger strengths of association with survival using the MOSTWAS model than the local-only model (**Supplemental Figure S6**). QQ-plots for TWAS *Z*-statistics (**Supplemental Figure S7**) and *P*-values (**Supplemental Figure S8**) showed earlier departure from the null using local-only models. These results in TCGA-BRCA demonstrated the improved transcriptomic prediction and power to detect gene-trait associations using MOSTWAS over local-only modeling.

#### Functional hypothesis generation with MOSTWAS

We next conducted TWAS for breast cancer-specific survival using all genes with significant germline heritability at *P* < 0.05 with the most predictive MOSTWAS model (i.e. MeTWAS or DePMA model with the larger cross-validation *R*^2^ greater than 0.01). We identified 21 survival-associated loci at Benjamini-Hochberg FDR-adjusted *P* < 0.05. Of these 21 loci, 11 persisted when subjected to permutation testing at a significance threshold of FDR-adjusted *P* < 0.05 (colored red in **Figure 4D** and **Supplemental Table S4**).

An advantage of MOSTWAS is its ability to aid in functional hypothesis generation for mechanistic follow-up studies. The distal-SNP added-last test allows for identification of genes where trait association from distal variation is significant, above and beyond the contribution of the local component. For 8 of the TWAS-associated 11 loci, at FDR-adjusted *P* < 0.05, we found significant distal variation added-last associations (see **Supplemental Methods** and **Supplemental Table S4**), suggesting that distal variation may contribute to the gene-trait associations. All 8 of these loci showed distal association with the gene of interest mediated through a set of four transcription factors (*NAA50*, *ATP6V1A*, *ROCK2*, *USF3*), all highly interconnected within the MAPK pathway, known to be involved in breast cancer proliferation [38–43]. These regulatory sites serve as an example of how distal genomic regions can be prioritized for functional follow-up studies to elucidate the mechanisms underlying the SNP-gene-trait associations. These results showed the strength of MOSTWAS to detect and prioritize gene-trait associations that are influenced by distal variation and to aid in generating functional hypotheses for these distal relationships.

### Comparison of computation time

To assess the difference in computational burden between local-only, MeTWAS, and DePMA modeling, we randomly selected a set of 50 genes that are heritable across all three models from TCGA-BRCA and computed per-gene time for fitting models using a 24-core, 3.0 GHz processor. We found that MeTWAS (average of 225 seconds per gene) and DePMA (average 312 seconds per gene) took approximately 6-10 times longer to fit than a traditional local-only model (average 36 seconds), as shown in **Supplemental Figure S9**. Model-fitting here includes heritability estimation, estimating the SNP-expression weights, and cross-validation. We have implemented parallelized methods to train an expression model for a single gene in MOSTWAS. We also recommend fitting an entire set of genes from an RNA-seq panel via a batch computing approach [57–59]. Using a parallel implementation with 5 cores and batch computing, we trained MOSTWAS expression models for 15,568 genes from TCGA-BRCA in approximately 28 hours.

## DISCUSSION

Through simulation analysis and real applications using two datasets [23,24], we demonstrated that multi-omic methods that prioritize distal variation in TWAS have higher predictive performance and power to detect tissue-specific gene-trait associations [9,13,60], especially when distal variation contributes substantially to trait heritability. We proposed two methods (MeTWAS and DePMA) for identifying and including distal genetic variants in gene expression prediction models. We have provided implementations of these methods in MOSTWAS (Multi-Omic Strategies for Transcriptome-Wide Association Studies) R package, available freely on GitHub. MOSTWAS contains functions to train expression models with both MeTWAS and DePMA and outputs models with 5-fold cross-validation *R*^2^ ≥ 0.01 and significant non-zero germline heritability. The package also contains functions and documentation for simulation analysis [61], the weighted burden test for gene-trait associations [28] and follow-up permutation [4] and distal-SNPs added-last tests for TWAS using GWAS summary statistics. We also provide guidelines for parallelization to distribute computational across cores.

Not only does MOSTWAS improve transcriptomic imputation both in- and out-of-sample, it also provides a test for the identification of heritable mediators that affect eventual transcription of the gene of interest. These identified mediators can provide insight into the underlying mechanisms for SNP-gene-trait associations to improve detection of gene-trait associations and to prioritize biological units for functional follow-up studies. TWAS using MOSTWAS models was able to recapitulate 5 out of 14 known Alzheimer’s disease risk loci in IGAP GWAS summary statistics [49], which were not recoverable with local-only models. We showed the utility of the distal-SNPs added-last test to prioritize significant distal SNP-gene-trait associations for follow-up mechanistic studies, which could not be identified using traditional local-only TWAS. In PGC GWAS summary-level data for major depressive disorder [54], we found 102 risk loci, 7 of which were replicated in independent GWAX summary statistics from the UK Biobank [55]. Three of these seven loci (*SYT1*, *CACNA2D3*, *ADAD2*) encode important proteins involved in synaptic transmission in the brain and RNA editing. Studies have shown that variation at these loci may lead to loss of function at synapses and RNA editing that lead to psychiatric disorders [65–69]. Using MOSTWAS and iCOGs summary-level GWAS statistics for breast cancer-specific survival [34], we identified 11 survival-associated loci that are enriched for p53 binding and oxidoreductase activity pathways [62,63]. These loci include two genes (*MAP3K6* and *MAP4K5*) encoding mitogen-activated protein kinases, which are signaling transduction molecules involved in the progression of aggressive breast cancer hormone subtypes [64]. None of the risk- or survival-associated loci identified by MOSTWAS were detected using local-only models.

A considerable limitation of MOSTWAS is the increased computational burden over local-only modeling, especially in DePMA’s permutation-based mediation analysis for multiple genome-wide mediators. By making some standard distributional assumptions on the SNP-mediator effect size and mediator-gene effect size vectors (e.g. effect sizes following a correlated multivariate Normal distribution), we believe a Monte-Carlo resampling method to estimate the null distribution of the product of these two effect size vectors may decrease computational time without significant loss in statistical power [70]. Nevertheless, we believe that MOSTWAS’s gain in predictive performance and power to detect gene-trait associations outweighs the added computational cost. Another concern with the inclusion of distal variants is that RNA-sequencing alignment errors can lead to false positives in distal-eQTL detection [71], and in turn, bias the mediation modeling. Cross-mapping estimation, as described by Saha *et al*, can be used to flag potential false positive distal-QTLs that are detected in the first step of MeTWAS and DePMA. Another limitation of MOSTWAS is the general lack of rich multi-omic panels, like ROS/MAP and TCGA-BRCA, that provide a large set of mediating biomarkers that may be mechanistically involved in gene regulation. However, the two-step regression framework outlined in MeTWAS allows for importing mediator intensity models trained in other cohorts to estimate the germline portion of total gene expression from distal variants. Importing mediator models from an external cohort can also reduce the testing burden in the preliminary QTL analysis in MeTWAS and DePMA.

MOSTWAS provides a user-friendly and intuitive tool that extends transcriptomic imputation and association studies to include distal regulatory genetic variants. We demonstrate that the methods in MOSTWAS based on two-step regression and mediation analysis generally out-perform local-only models in both transcriptomic prediction and TWAS power without signs of inflated false positive rates, though at the cost of longer computation time. MOSTWAS enables users to utilize rich reference multi-omic datasets for enhanced gene mapping to better understand the genetic etiology of polygenic traits and diseases with more direct insight into functional follow-up studies.

## MATERIALS AND METHODS

We first outline the two methods proposed in this work: (1) mediator-enriched TWAS (MeTWAS) and (2) distal-eQTL prioritization via mediation analysis (DePMA). MeTWAS and DePMA are combined in the MOSTWAS R package, available at www.github.com/bhattacharya-a-bt/MOSTWAS. Full mathematical details are provided in **Supplemental Methods**.

### Transcriptomic prediction using MeTWAS

Across all samples in the training dataset and for a single gene of interest, MeTWAS, an adaptation of two-step regression, takes in a vector of gene expression, the matrix of genotype dosages local to the gene of interest (default of 0.5 Megabases around the gene), and a set of mediating biomarkers that are estimated to be significantly associated with the expression of the gene of interest through a QTL analysis. In accord with previous studies that use penalized regression methods [35,72,73], we only select the most significant gene-associated mediators as adding too many potentially redundant features often leads to poorer predictive performance. This feature selections also limits computational time. Through simulations, we observed that including all SNPs local to the mediators results in lower predictive *R*^2^ compared to the two-step regression method in MeTWAS (**Supplemental Figure S10**), and the discrepancy between these methods is larger in practice (results not shown). These mediating biomarkers can be DNA methylation sties, microRNAs, transcription factors, or any molecular feature that may be genetically heritable and affect transcription.

Transcriptome prediction in MeTWAS draws from two-step regression, as summarized in **Supplemental Figure S1**. Using the genotype local to these mediators, MeTWAS first trains a predictive model for their intensities (i.e. expression, methylation, etc.) using either elastic net [26] or linear mixed modeling [27]. In practice, we found that a simpler, one-step procedure of including all variants local to both the gene and to potential mediators led to the distal SNP effects being estimated as zero during the regularization process, even in simulations when the true distal SNP effects were nonzero (see **Methods**). We then use these predictive models to estimate the genetically regulated intensity (GRIn) of each mediator in the training set, through cross-validation. The GRIns for each mediator is then included in a matrix of fixed effects. The effect sizes of the GRIns on the expression of the gene of interest are estimated using ordinary least squares regression, and then the expression vector is residualized for these effect sizes. Effect sizes of variants local to the gene of interest are then estimated using elastic net or linear mixed modeling [26,27] on the residualized gene expression quantity. Details are provided in **Supplemental Methods**.

### Transcriptomic prediction using DePMA

Expression prediction in DePMA hinges on prioritizing distal-eSNPs via mediation analysis for inclusion in the final DePMA predictive model, adopting methods from previous studies [11,12,14]. A multi-omic dataset with gene expression, SNP dosages, and potential mediators is first split into training-testing subsets. Based on the minor allele frequencies of SNPs and total sample size, we recommend a low number of splits (less than 5).

In the training set, we identify mediation test triplets that consist of (1) a gene of interest, (2) a distal-eSNP associated with the expression of the gene (default of *P* < 10^−6^), and (3) a set of mediating biomarkers local to and associated with the distal-eSNP (default of FDR-adjusted *P* < 0.05). We estimate the total indirect mediation effect (TME) of the distal-eSNP on the gene of interest mediated through the set of these mediators, as defined by Sobel [74]. We assess the magnitude of this indirect effect using a two-sided permutation test to obtain a permutation *P*-value, as more direct methods of computing standard errors for the estimated TME are often biased [14,75]. We also provide an option to estimate an asymptotic approximation to the standard error of the TME and conduct a Wald-type test. This asymptotic option is significantly faster at the cost of inflated false positives (see **Supplemental Methods** and **Supplemental Figure S11**). Distal-eSNPs with significantly large absolute TMEs are included with the local SNPs for the gene of interest in a predictive model, fit using elastic net or linear mixed modeling [26,27]. These SNP effect sizes can then be exported for imputation in external GWAS cohorts. Details are provided in **Supplemental Methods**.

### Transcriptomic imputation with MOSTWAS

In an external GWAS panel, if individual level genotypes are available, we construct the mediator-enriched genetically regulated expression (GReX) of the gene of interest by multiplying the genotypes in the GWAS panel by the effect sizes estimated in a MOSTWAS model. This GReX quantity represents the component of total expression that is attributed to germline genetics and can be used in downstream TWAS to detect gene-trait associations.

### Tests of association

If individual level genotypes are not available, then the weighted burden *Z*-test, proposed by Pasaniuc *et al* and employed in FUSION [4,28], can be employed and applied to summary statistics. Briefly, the test statistic is a linear combination of the *Z*-scores corresponding to the SNPs included in the MOSTWAS model for a gene of interest, where each individual GWAS *Z*-score is weighted by the corresponding MOSTWAS effect size. The covariance matrix for this weighted burden test statistic is estimated from the linkage disequilibrium between SNPs in the eQTL panel or some publicly available ancestry-matched reference panels. This weighted burden test statistic is compared to the standard Normal distribution for inference.

We implement a permutation test, conditioning on the GWAS effect sizes to assess whether the same distribution SNP effect sizes could yield a significant association by chance [4]. We permute the effect sizes 1,000 times without replacement and recompute the weighted burden test statistic to generate permutation null distribution. This permutation test is only conducted for overall associations at a user-defined significance threshold (default to FDR-adjusted *P* < 0.05).

Lastly, we also implement a test to assess the information added from distal-eSNPs in the weighted burden test beyond what we find from local SNPs. This test is analogous to a group added-last test in regression analysis, applied here to GWAS summary statistics. Formally, we test whether the weighted burden test statistic for the distal-SNPs is significantly non-zero given the observed weighted burden test statistic for the local-SNPs. We draw conclusions from the assumption that these two weighted burden test statistics follow bivariate Normal distribution. Full details and derivations are given in **Supplemental Methods**.

### Simulation framework

We first conducted simulations to assess the predictive capability and power to detect gene-trait associations under various settings for phenotype heritability, local and distal heritability of expression, and proportion of causal local and distal SNPs. We considered two scenarios: We considered two scenarios for each set of simulation parameters: (1) an ideal case where the leveraged associated between the distal-SNP and gene of interest exists in both the reference and imputation panel, and (2) a “null” case where the leveraged association between the distal-SNP and the gene of interest exists in the reference panel but does not contribute phenotype heritability in the imputation panel.

Using genetic data from TCGA-BRCA as a reference, we used SNPs local to the gene *ESR1* (Chromosome 6) to generate local eQTLs and SNPs local to *FOXA1* (Chromosome 14) to generate distal-eQTLs for a 400-sample eQTL reference panel and 1,500-sample GWAS imputation panel, as in Mancuso *et al*’s *twas_sim* protocol [61]. We computed the adjusted predictive *R*^2^ in the reference panel for the trained MeTWAS and DePMA models and tested the gene-trait association in the GWAS panel using the weighted burden test. The association study power was defined as the proportion of gene-trait associations with *P* < 2.5 × 10^−6^, the Bonferroni-corrected significance threshold for testing 20,000 independent genes. With these simulated datasets, we also assessed the power of the distal added-last test by computing the proportion of significant distal associations, conditional on the local association at FDR-adjusted *P* < 0.05. Full details are provided in **Supplemental Methods**.

### Data acquisition

#### Multi-omic data from ROS/MAP

We retrieved imputed genotype, RNA expression, miRNA expression, and DNA methylation data from The Religious Orders Study and Memory and Aging Project (ROS/MAP) Study for samples derived from human pre-frontal cortex [23,82,83]. We excluded variants (1) with a minor allele frequency of less than 1% based on genotype dosage, (2) that deviated significantly from Hardy-Weinberg equilibrium (*P* < 10^−8^) using appropriate functions in PLINK v1.90b3 [79,80], and (3) located on sex chromosomes. Final ROS/MAP genotype data was coded as dosages, with reference and alternative allele coding as in dbSNP. We intersected to the subset of samples assayed for genotype (at 4,141,537 variants), RNA-seq (15,857 genes), miRNA-seq (247 miRNAs), and DNA methylation (391,626 CpG sites), resulting in a total of 370 samples. Again, we only considered the autosome in our analyses. We adjusted gene and miRNA expression and DNA methylation by relevant covariates (10 principal components of the genotype age at death, sex, and smoking status).

#### Multi-omic data from TCGA-BRCA

We retrieved genotype, RNA expression, miRNA expression, and DNA methylation data for breast cancer indications in The Cancer Genome Atlas (TCGA). Birdseed genotype files of 914 subjects were downloaded from the Genome Data Commons (GDC) legacy (GRCh37/hg19) archive. Genotype files were merged into a single binary PLINK file format (BED/FAM/BIM) and imputed using the October 2014 (v.3) release of the 1000 Genomes Project dataset as a reference panel in the standard two-stage imputation approach, using SHAPEIT v2.87 for phasing and IMPUTE v2.3.2 for imputation [76–78]. We excluded variants (1) with a minor allele frequency of less than 1% based on genotype dosage, (2) that deviated significantly from Hardy-Weinberg equilibrium (*P* < 10^−8^) using appropriate functions in PLINK v1.90b3 [79,80], and (3) located on sex chromosomes. Final TCGA genotype data was coded as dosages, with reference and alternative allele coding as in dbSNP.

TCGA level-3 normalized RNA-seq expression data, miRNA-seq expression data, and DNA methylation data collected on Illumina Infinium HumanMethylation450 BeadChip were downloaded from the Broad Institute’s GDAC Firehose (2016/1/28 analysis archive) via FireBrowse [24,81]. We intersected to the subset of samples assayed for genotype (4,564,962 variants), RNA-seq (15,568 genes), miRNA-seq (1,046 miRNAs), and DNA methylation (485,578 CpG sites), resulting in a total of 563 samples. We only considered the autosome in our analyses. We adjusted gene and miRNA expression and DNA methylation by relevant covariates (10 genotype principal components, tumor stage at diagnosis, and age).

#### Summary statistics for downstream association studies

We conducted TWAS association tests using relevant GWAS summary statistics for breast cancer-specific survival, risk of late-onset Alzheimer’s disease, and risk of major depressive disorder. We downloaded iCOGs GWAS summary statistics for breast cancer-specific survival for women of European ancestry [34]. All studies and funders as listed in Michailidou *et al* [32,33] and in Guo *et al* [34] are acknowledged for their contributions. Furthermore, we downloaded GWAS summary statistics for risk of late-onset Alzheimer’s disease from the International Genomics of Alzheimer’s Project (IGAP) [49].

We also downloaded GWAS and genome-wide association by proxy (GWAX) summary statistics for risk of major depressive disorder (MDD) from the Psychiatric Genomics Consortium [54] and the UK Biobank [55], respectively. IGAP is a large two-stage study based on GWAS on individuals of European ancestry. In stage 1, IGAP used genotyped and imputed data on 7,055,881 single nucleotide polymorphisms (SNPs) to meta-analyze four previously-published GWAS datasets consisting of 17,008 Alzheimer’s disease cases and 37,154 controls (The European Alzheimer’s disease Initiative – EADI, the Alzheimer Disease Genetics Consortium – ADGC, The Cohorts for Heart and Aging Research in Genomic Epidemiology consortium – CHARGE, The Genetic and Environmental Risk in AD consortium – GERAD). In stage 2, 11,632 SNPs were genotyped and tested for association in an independent set of 8,572 Alzheimer’s disease cases and 11,312 controls. Finally, a meta-analysis was performed combining results from stages 1 and 2.

### Model training and association testing in ROS/MAP and TCGA-BRCA

Using both ROS/MAP and TCGA-BRCA multi-omic data, we first identified associations between SNPs and mediators (transcription factor genes, miRNAs, and CpG methylation sites), mediators and gene expression, and SNPs and gene expression using MatrixEQTL [84]. These QTL analyses were adjusted for 10 genotype principal components to account for population stratification, along with other relevant covariates (age, sex, and smoking status for ROS/MAP; tumor stage and age for TCGA-BRCA). For MeTWAS modeling, we considered the top 5 mediators associated with the gene of interest, assessed by the smallest FDR-adjusted *P* < 0.05. For DePMA models, we considered all distal-SNPs associated with gene expression at raw *P* < 10^−6^ and any local mediators at FDR-adjusted *P* < 0.05. Local windows for all models were set to 0.5 Mb. For association testing, we considered only genes with significant non-zero estimated total heritability by GCTA-LDMS [85] and cross-validation adjusted *R*^2^ ≥ 0 across 5 folds. The MeTWAS or DePMA model with larger cross-validation *R*^2^ was considered as the final MOSTWAS model for each gene. All other modeling options in MeTWAS and DePMA were set to the defaults provided by the MOSTWAS package.

Using ROS/MAP models, we first conducted TWAS burden testing in GWAS summary statistics for late-onset Alzheimer’s disease risk from IGAP, prioritized 14 known risk loci identified from literature [49–52]. We subjected TWAS-identified loci at FDR-adjusted *P* < 0.05 to permutation testing, and any loci that persisted past permutation testing to distal variation added-last testing. We similarly conducted TWAS for risk of major depressive disorder (MDD) using GWAS summary statistics from PGC (excluding data from 23andMe and the UK Biobank) with the necessary follow-up tests. For any TWAS-identified loci that persisted permutation in PGC, we further conducted TWAS in GWAX summary statistics for MDD risk in the UK Biobank [55] for replication.

Using TCGA-BRCA models, we conducted TWAS burden testing [4,28] in iCOGs GWAS summary statistics for breast cancer-specific survival in a cohort of women of European ancestry. We subjected TWAS-identified loci at Benjamini-Hochberg [37] FDR-adjusted *P* < 0.05 to permutation testing, and any locus that persisted past permutation testing to distal variation added-last testing.

## ACKNOWLEDGEMENTS

We thank Melissa Troester, Colin Begg, Terry Furey, Michael Gandal, Sasha Gusev, Karen Mohlke, Brandon Pierce, Bogdan Pasaniuc, Hudson Santos, and Jason Stein for engaging conversation and guidance during the research process.

We thank the International Genomics of Alzheimer’s Project (IGAP) for providing summary results data for these analyses. The investigators within IGAP contributed to the design and implementation of IGAP and/or provided data but did not participate in analysis or writing of this report. IGAP was made possible by the generous participation of the control subjects, the patients, and their families. The iSelect chips were funded by the French National Foundation on Alzheimer’s disease and related disorders. EADI was supported by the LABEX (laboratory of excellence program investment for the future) DISTALZ grant, Inserm, Institut Pasteur de Lille, Université de Lille 2 and the Lille University Hospital. GERAD was supported by the Medical Research Council (Grant n° 503480), Alzheimer’s Research UK (Grant n° 503176), the Wellcome Trust (Grant n° 082604/2/07/Z) and German Federal Ministry of Education and Research (BMBF): Competence Network Dementia (CND) grant n° 01GI0102, 01GI0711, 01GI0420. CHARGE was partly supported by the NIH/NIA grant R01 AG033193 and the NIA AG081220 and AGES contract N01–AG–12100, the NHLBI grant R01 HL105756, the Icelandic Heart Association, and the Erasmus Medical Center and Erasmus University. ADGC was supported by the NIH/NIA grants: U01 AG032984, U24 AG021886, U01 AG016976, and the Alzheimer’s Association grant ADGC–10–196728.

We also thank the Psychiatric Genomics Consortium for their publicly available GWAS summary statistics (https://www.med.unc.edu/pgc/) and the Pickrell Lab at the New York Genome Center for their GWAS browser and GWAX summary statistics (http://gwas-browser.nygenome.org/downloads/gwas-browser/).

Funding for BCAC and iCOGS came from: Cancer Research UK [grant numbers C1287/A16563, C1287/A10118, C1287/A10710, C12292/A11174, C1281/A12014, C5047/A8384, C5047/A15007, C5047/A10692, C8197/A16565], the European Union’s Horizon 2020 Research and Innovation Programme (grant numbers 634935 and 633784 for BRIDGES and B-CAST respectively), the European Community’s Seventh Framework Programme under grant agreement n° 223175 [HEALTHF2-2009-223175] (COGS), the National Institutes of Health [CA128978] and Post-Cancer GWAS initiative [1U19 CA148537, 1U19 CA148065-01 (DRIVE) and 1U19 CA148112 – the GAME-ON initiative], the Department of Defence [W81XWH-10-1-0341], and the Canadian Institutes of Health Research CIHR) for the CIHR Team in Familial Risks of Breast Cancer [grant PSR-SIIRI-701]. All studies and funders as listed in Michailidou K *et al* (2013 and 2015) and in Guo Q et al (2015) are acknowledged for their contributions.

## SUPPORTING INFORMATION

Document S1: Supplemental Methods (SuppMethods.pdf)

Document S2: Supplemental Figures (Supplemental Figures S1-S11; SuppFigs.pdf)

Document S3: Supplemental Table (Supplemental Tables S1-S4; SuppTables.pdf)

## DATA AVAILABILITY

MOSTWAS software, https://github.com/bhattacharya-a-bt/MOSTWAS

Models and full results, https://zenodo.org/record/3755919 [30]

TCGA GDC Legacy Archive, https://portal.gdc.cancer.gov/legacy-archive

GDAC Firehose Browser, https://gdac.broadinstitute.org

ROS/MAP data, https://www.synapse.org/#!Synapse:syn3219045

iCOGS GWAS Summary Statistics, http://bcac.ccge.medschl.cam.ac.uk/bcacdata/icogs-complete-summary-results

IGAP Late-onset Alzheimer’s Disease Risk GWAS Summary Statistics, http://web.pasteur-lille.fr/en/recherche/u744/igap/igap_download.php

PGC Major Depressive Disorder GWAS Summary Statistics, https://www.med.unc.edu/pgc/download-results/mdd/

UKBB Major Depressive Disorder GWAX Summary Statistics, http://gwas-browser.nygenome.org/downloads/gwas-browser/

## AUTHOR CONTRIBUTIONS

Conceptualization: AB, MIL; Data Curation: AB; Formal Analysis: AB; Funding Acquisition: MIL, YL; Methodology: AB, MIL; Software: AB; Supervision: MIL; Validation and visualization: AB, MIL, YL; Writing – Original Draft Preparation: AB; Writing – Review and Editing: MIL, YL

