## Supplemental Figures for "MOSTWAS: Multi-Omic Strategies for Transcriptome-Wide Association Studies"

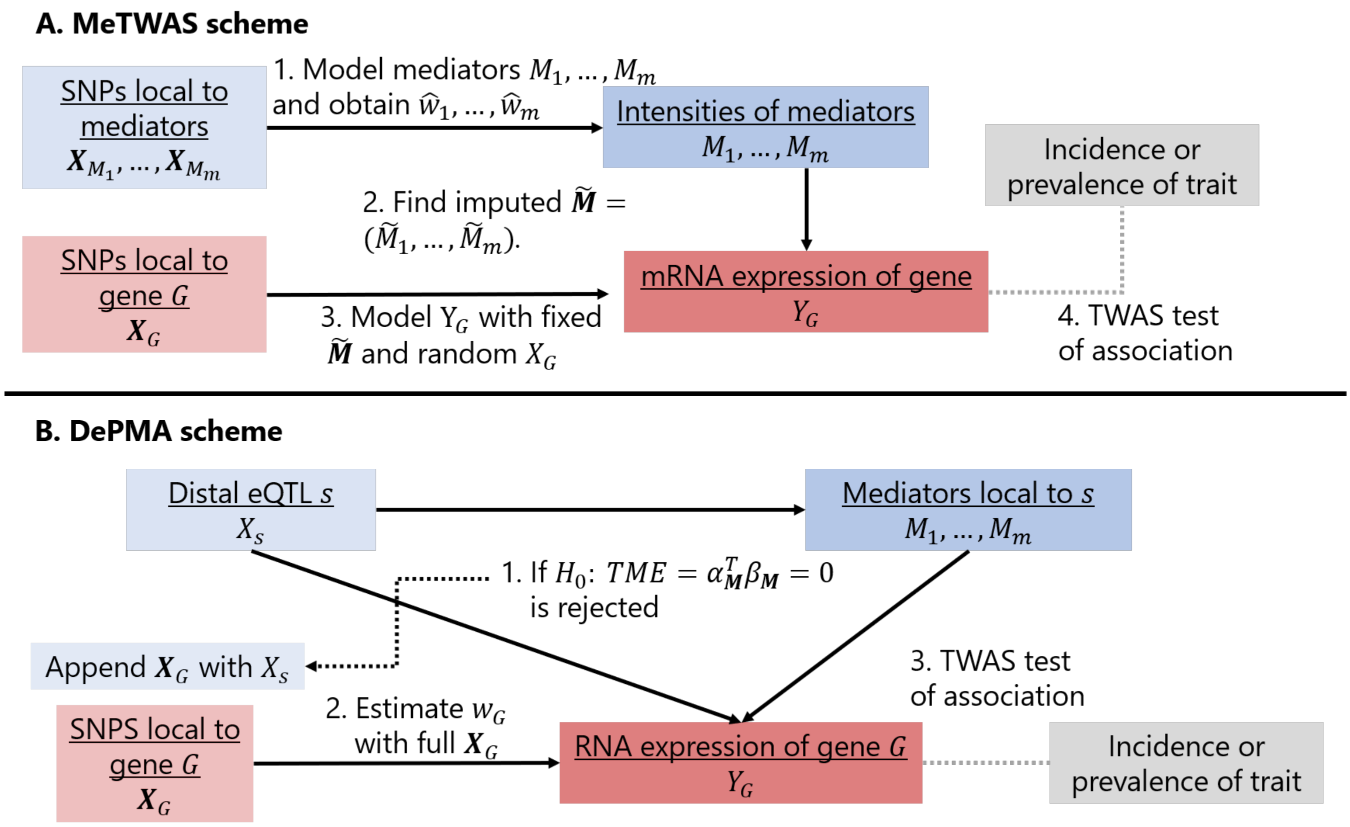


Figure S1: *Algorithmic details for mediator-enriched TWAS (MeTWAS) and distal-eQTL prioritization via mediation analysis (DePMA).*

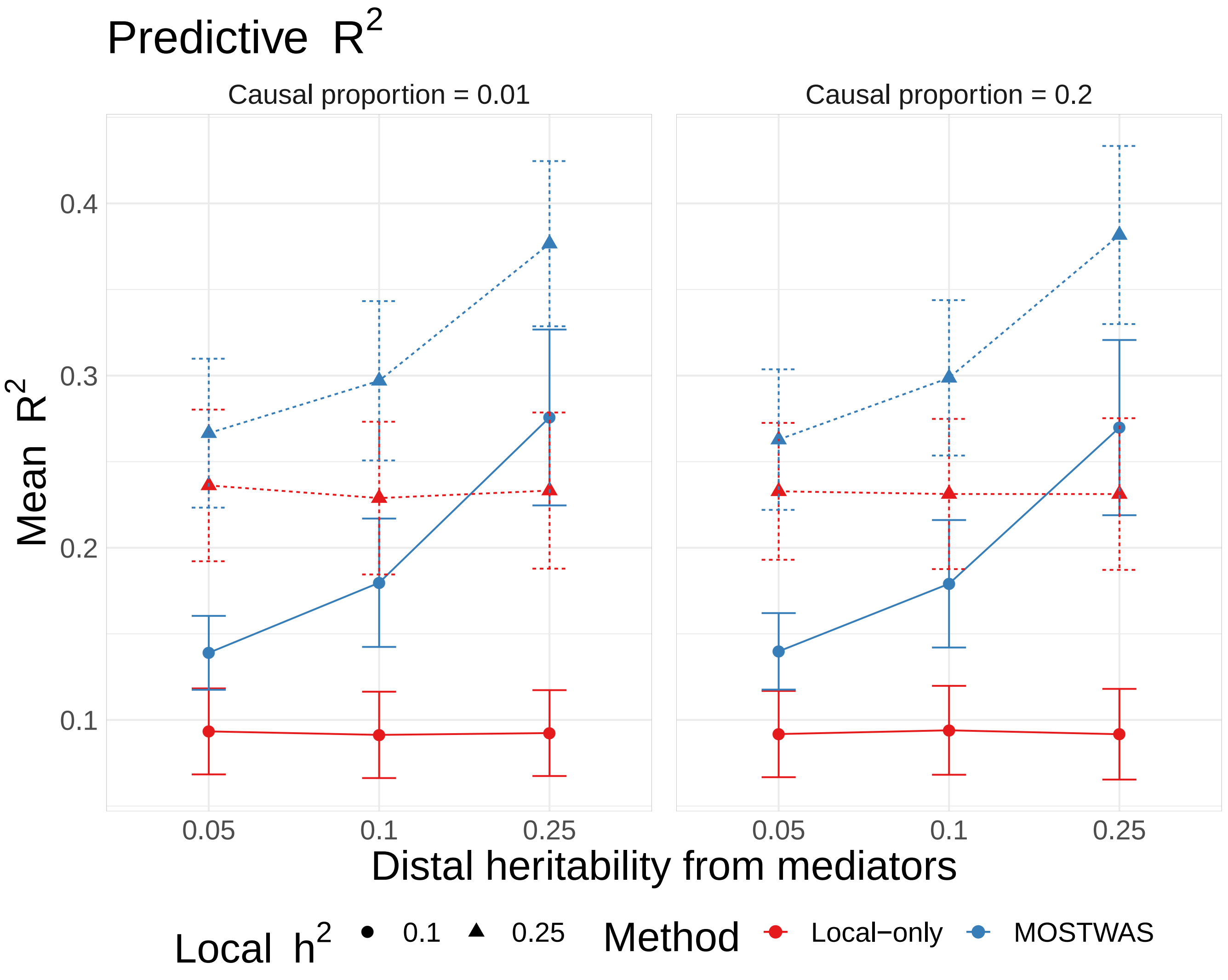


Figure S2: *Comparison of predictive $R^2$ in simulations.* Mean adjusted $R^2$ across various local and distal expression heritabilities, trait heritabilities, and causal proportions using local-only (red) and the best MOSTWAS (blue) models. The error bars reflect a width of 1 standard deviation of the 1,000 simulated adjusted $R^2$ values.

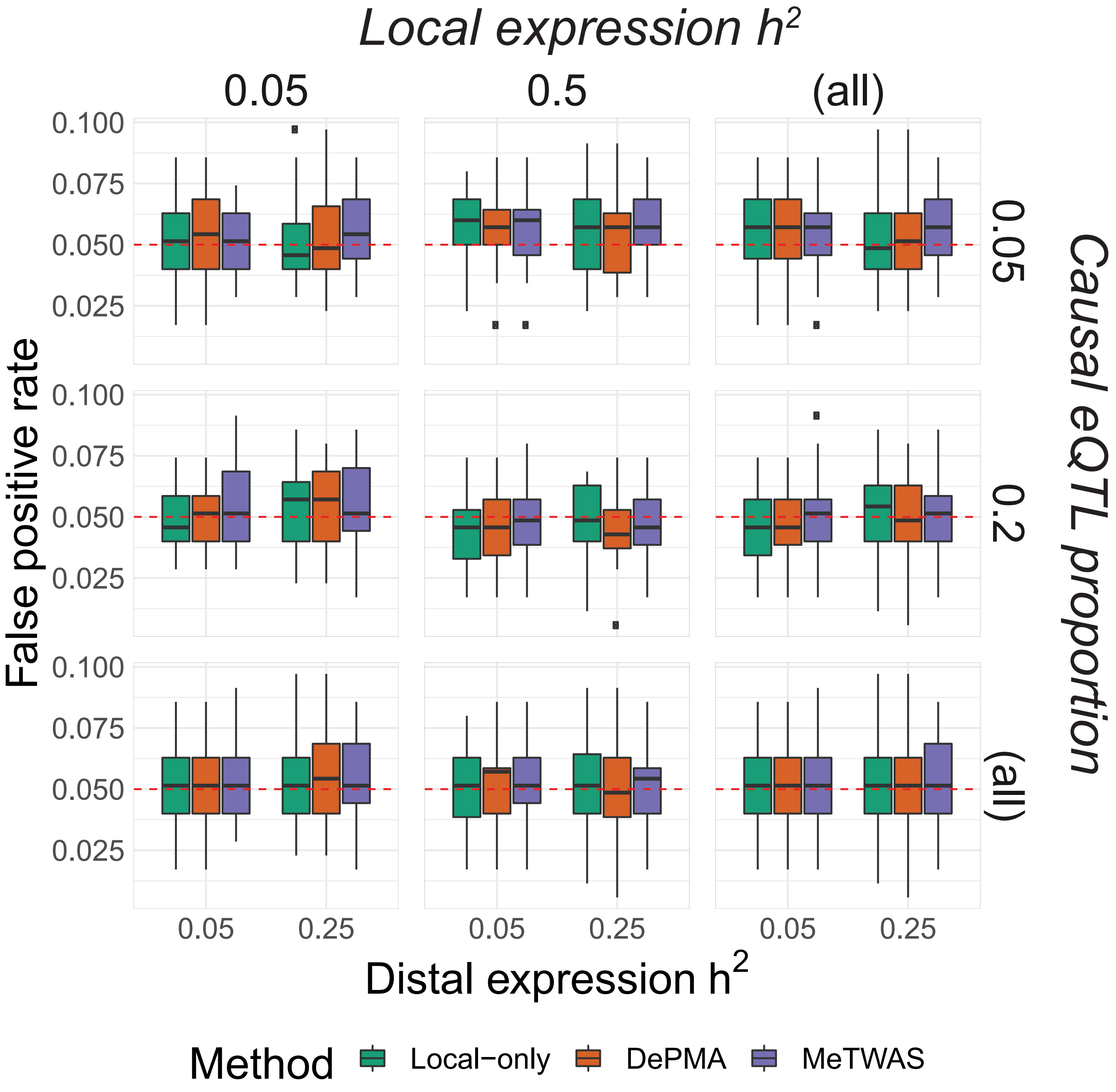


Figure S3: *Comparison of false positive rates in simulations.* Boxplots of false positive rate ($Y$-axis) across various distal expression heritability settings ($X$-axis), stratified by local expression heritability (horizontal) and causal eQTL proportion (vertically) and colored by the method used to test gene-trait associations. These boxplots are from 20 simulations of 1,000 permuted phenotype traits in the simulated GWAS. The red line provides a reference at a false positive rate of 0.05.

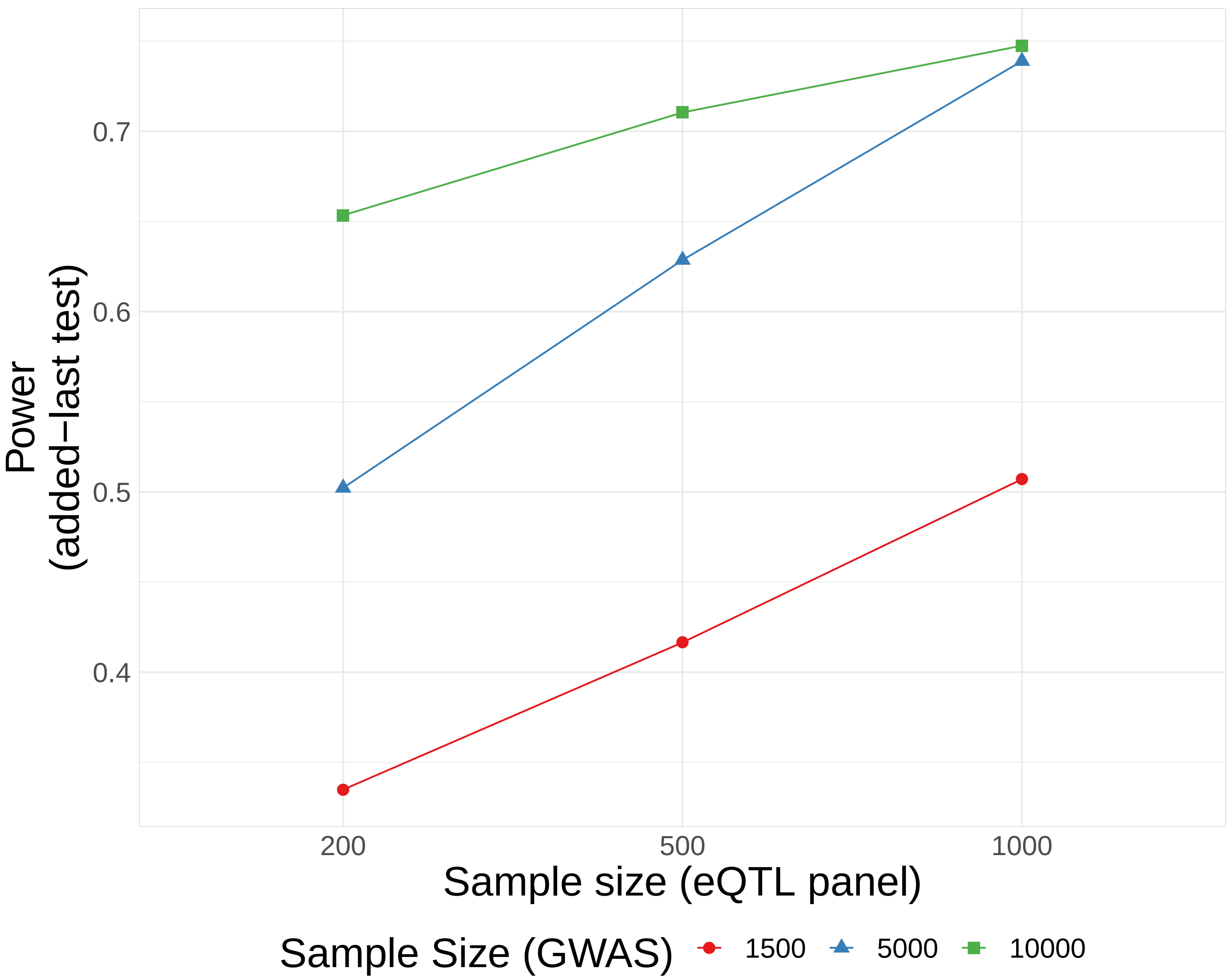


Figure S4: *Simulation analysis for the power of the distal variants added-last test.* Across various sample sizes for the eQTL reference ($X$-axis) panel and GWAS imputation panel (color), the power of the distal added-last test to detect a significant association with distal variants conditional on a significant local association at FDR-adjusted $P < 0.05$.

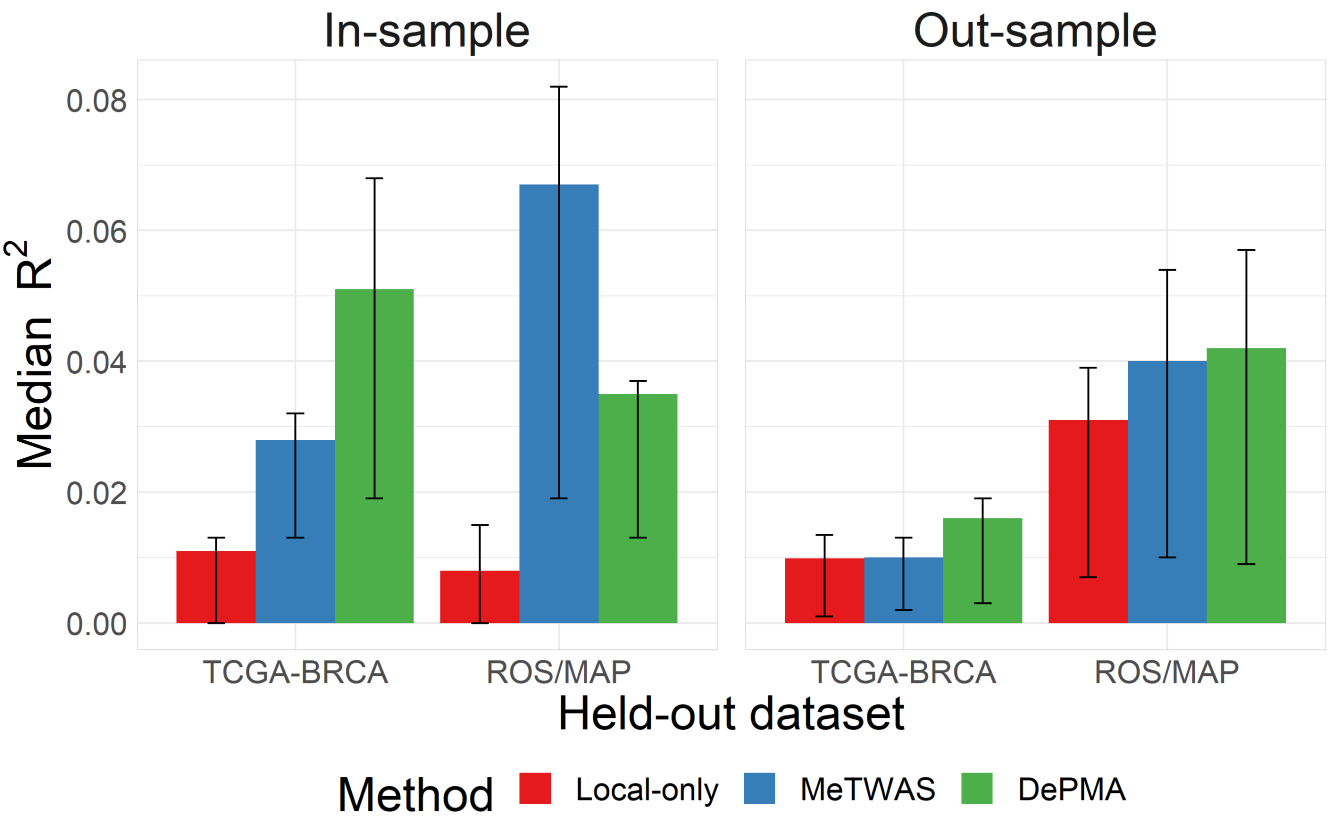


Figure S5: *Comparison of in- and out-sample predictive performance of local-only and MOSTWAS expression models.* Median predictive adjusted $R^2$ for in-sample (left) and out-sample (right) performance in TCGA-BRCA and ROS/MAP expression models using local-only (red), MeTWAS (blue), and DePMA (green) modelling. The interval provided shows the 25% and 75% quartiles. Only genes with significant $h^2$ at raw $P < 0.05$ are shown here.

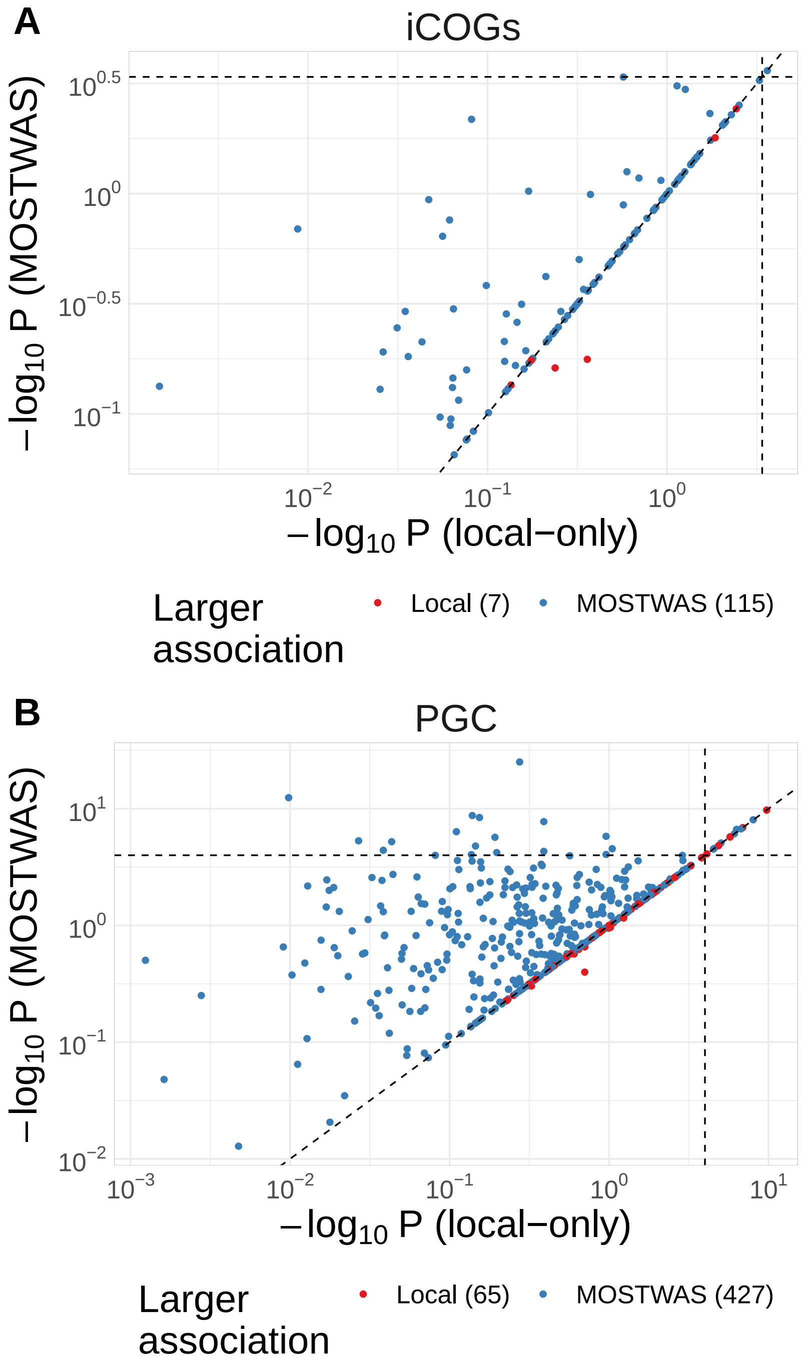


Figure S6: *Gene-trait associations in iCOGs and PGC using local-only and MOSTWAS models.* $-log_{10}P$-values of weighted burden gene-trait associations using iCOGs survival GWAS in European-ancestry women (left) and PGC MDD risk GWAS in predominantly European-ancestry patients (right) among genes that were predicted at cross-validation $R^2 \geq 0.01$ using both local-only and MOSTWAS models. The $X$- and $Y$-axes display the $-log_{10}P$-values for local-only and the best MOSTWAS model, respectively, on a log-scale. Points are colored red if $P$-value of association is less than or equal using the MOSTWAS model. The horizontal and vertical reference lines indicate overall Bonferroni-corrected significance thresholds ($4.1 \times 10^-4$ for iCOGs, $1.0 \times 10^-4$ for PGC.

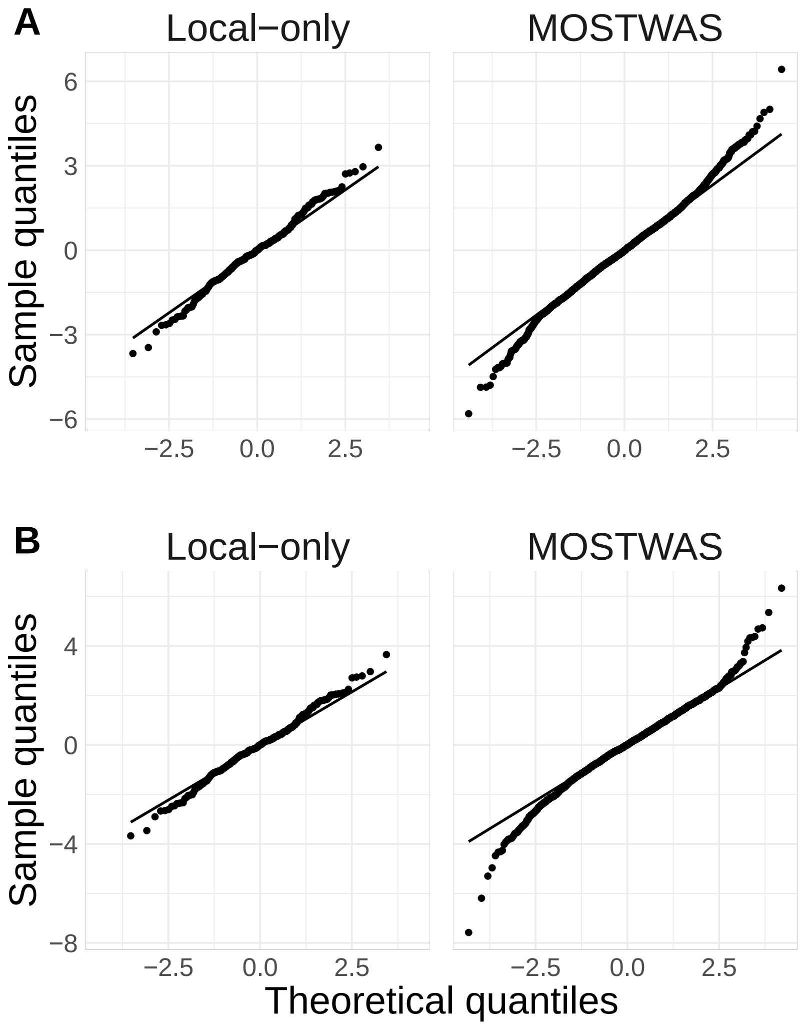


Figure S7: *Comparison of QQ-plots from TWAS associations.* QQ-plots of $Z$ scores from TWAS for breast cancer-specific survival in iCOGs (A) and MDD in PGC (B) with local-only models (left) and MOSTWAS (right)

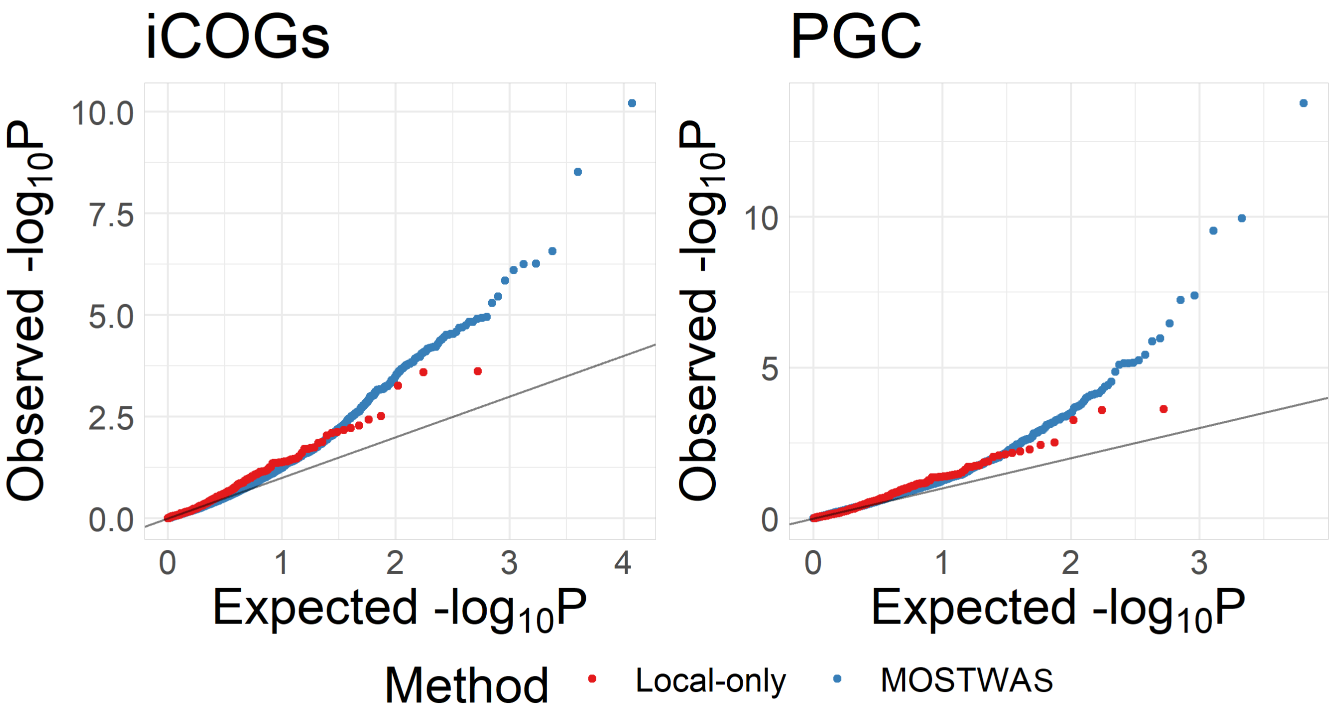


Figure S8: *Comparison of QQ-plots from TWAS associations.* QQ-plots of $-\log_{10}$ $P$-values from TWAS for breast cancer-specific survival in iCOGs (left) and MDD in PGC (right) with local-only models and MOSTWAS models.

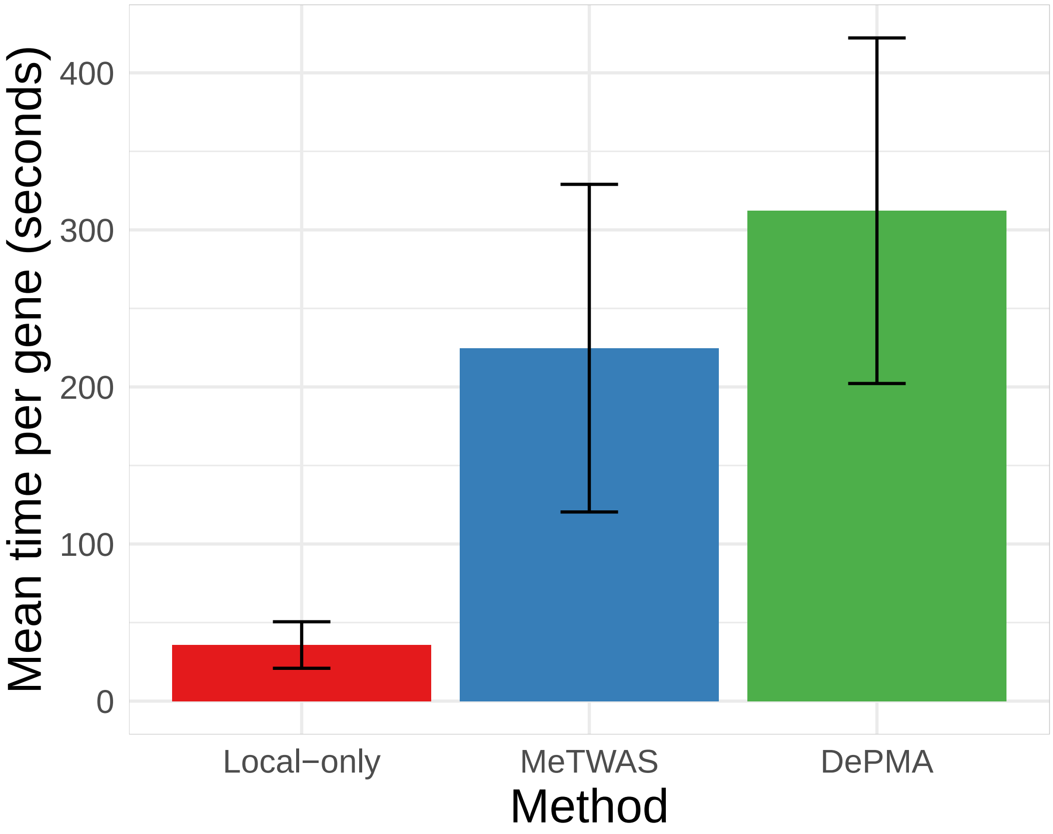


Figure S9: *Comparison of computation times between local-only and MOSTWAS modelling.* Mean and standard deviation of per-gene computation time across 50 randomly selected genes in TCGA-BRCA. Computations here were done with a 24-core, 3.0 GHz processor.

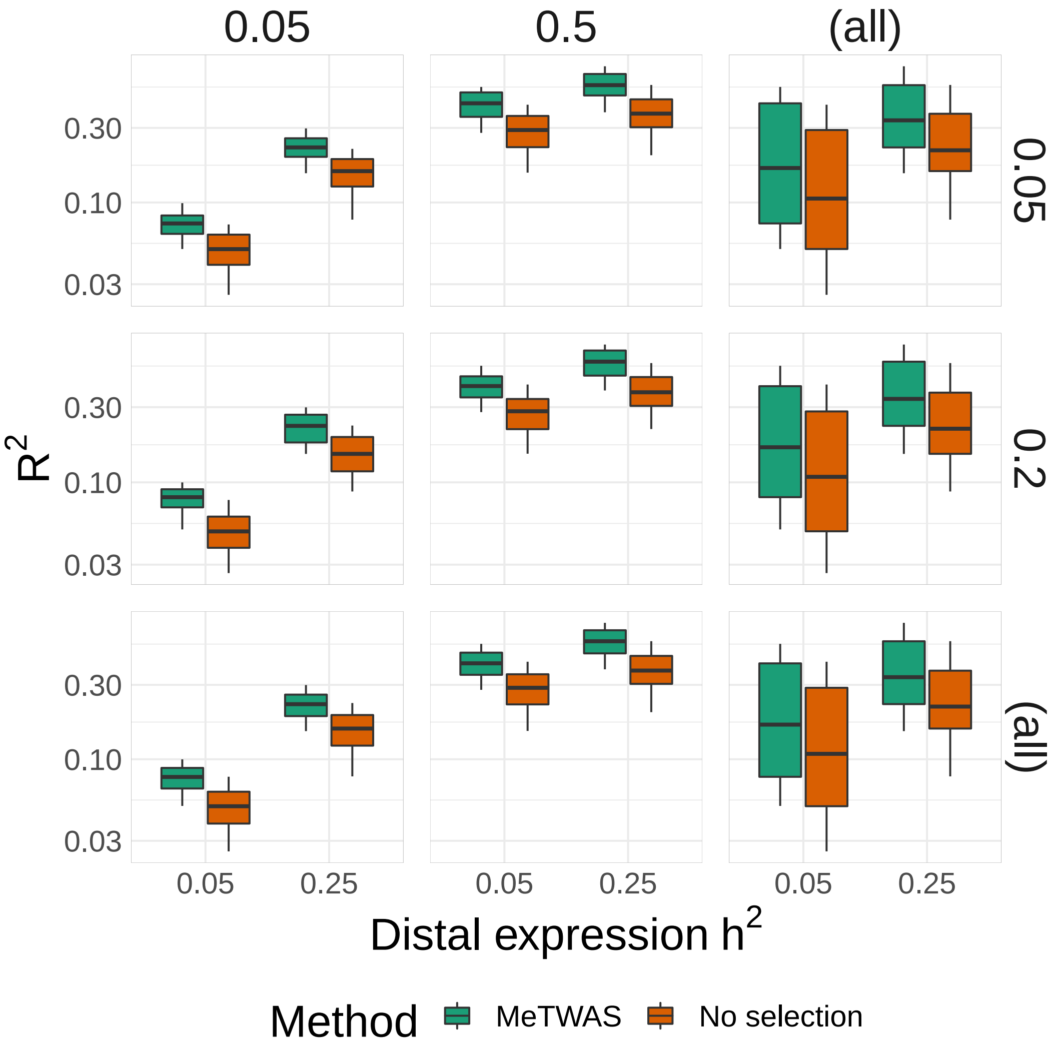


Figure S10: *Comparison of predictive ability of MeTWAS with one- and two-step regression.* Comparison of predictive performance ($Y$-axis) of two-step regression (labelled as MeTWAS) and one-step regression (labelled no selection) across 100 simulations across various causal proportions of eQTLs (vertically arranged), local expression heritability (horizontal), and distal expression heritability ($X$-axis).

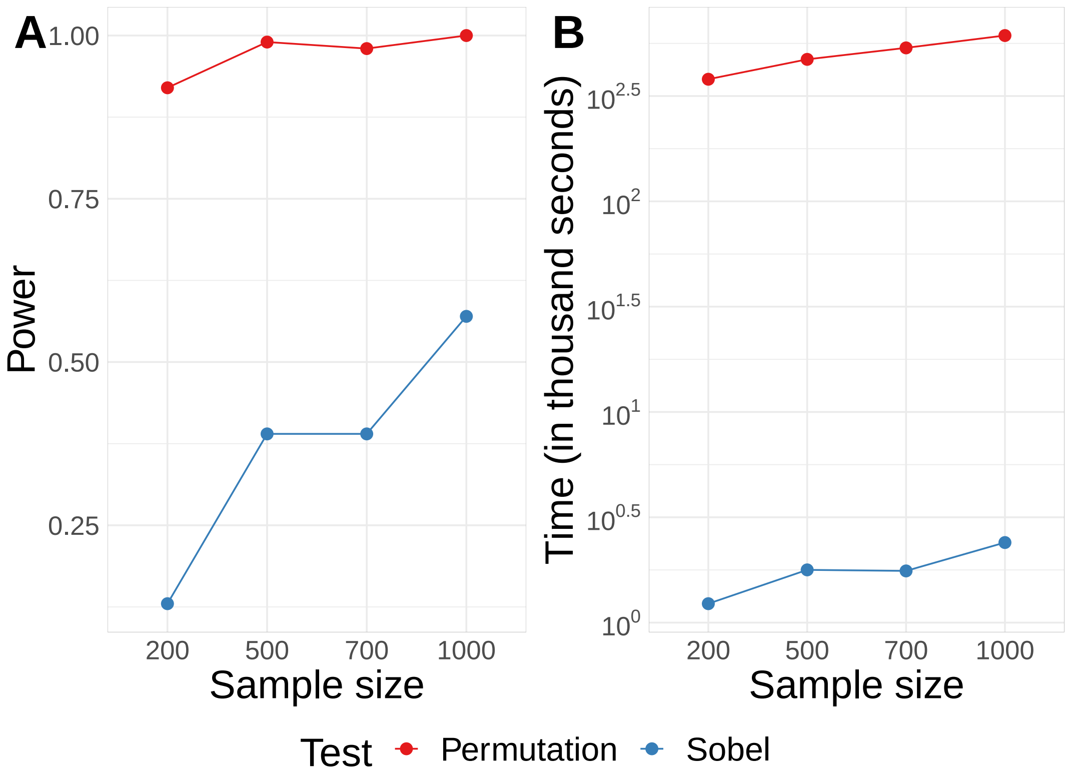


Figure S11: *Comparison of test power and computational speed of Sobel asymptotic and permutation tests of total mediation effect.* Power ($Y$-axis, left) and computational time ($Y$-axis, right) to detect a true large absolute total mediation effect and computation speed over various eQTL panel sample sizes ($X$-axis) in 10,000 simulations of mediation testing triplets.
