## Supplemental Methods for "MOSTWAS: Multi-Omic Strategies for Transcriptome-Wide Association Studies"

We first outline the two methods proposed in this work, (1) mediator-enriched transcriptome-wide prediction (MeTWAS) and (2) distal-eQTL prioritization via mediation analysis (DePMA). MeTWAS and DePMA are combined in the MOSTWAS R package, available freely at `www.github.com/bhattacharya-a-bt/MOSTWAS`.

### 1 Mediator-enriched TWAS (MeTWAS)

#### 1.1 Transcriptomic prediction using MeTWAS

Here, we present mediator-enriched TWAS, or MeTWAS, one of the two tools presented in the MOSTWAS R package. Across $n$ samples, consider the vector $Y_G$, the expression of a gene $G$ of interest, the matrix $\mathbf{X}_G$ of local SNPs dosages in a user-defined window around gene $G$ (default to 0.5 Megabases), and $m_G$ mediating biomarkers that we estimate to be significantly associated with the expression of gene $G$ via a relevant one-way test of association. These mediating biomarkers can be DNA methylation sites, microRNAs, transcription factors, or any molecular profile that may be genetically heritable and affect transcription. Accordingly, let the matrix $\mathbf{X}_{M_j}$ be the local-genotype dosages in a 500 kilobase window around mediator $j$, $1 \leq j \leq m_G$. Furthermore, let $M_j$ be the intensity of mediator $j$ (methylation $M$-value if $j$ is a CpG site, expression if $j$ is a miRNA or a gene, etc). Prior to any modeling, we scale $Y_G$ and all $M_j$, $1 \leq j \leq m_G$ to zero mean and unit variance. We also residualize $M_j$, $1 \leq j \leq m_G$ and $Y_G$ with the covariate matrix $\mathbf{X}_C$ to account for population stratification using principal components of the global genotype matrix and relevant clinical covariates to obtain $\tilde{M}_j$, $1 \leq j \leq m_G$ and $\tilde{Y}_G$.

Transcriptome prediction in MeTWAS draws from two-step regression, as summarized in **Supplemental Figure S1**. First, in the training set for a given training-test split, for $1 \leq j \leq m_G$, we model the residualized intensity $\tilde{M}_j$ of training-set specific mediator $j$ with the following additive model:

$$\tilde{M}_j = \mathbf{X}^{\dagger}_{M_j} w_j + \varepsilon_m, \tag{1}$$

where $w_j$ is the effect-sizes of the SNPs in $\mathbf{X}^{\dagger}_{M_j}$ on $\tilde{M}_j$ in the training set. As in traditional transcriptomic imputation models [1, 2], we find $\hat{w}_j$ using one of the two following methods with the largest predicted adjusted $R^2$: (1) elastic net regression with mixing parameter $\alpha = 0.5$ and $\lambda$ tuned over 5-fold cross validation using glmnet [3], or (2) linear mixed modeling assuming random effects for $\mathbf{X}_{M_j}$ using rrBLUP [4]. Only significantly heritable (default $P < 0.05$ for the likelihood ratio test) [5] and well-cross validated (default $R^2 \geq 0.01$) expression models are considered.

For all $j$, using these optimized predictive models for $M_j$ as denoted by $\hat{w}_{M_j}$, we estimate the genetically regulated intensity (GRIn) of the mediator $m_j$, denoted $\text{GRIn}_{m_j}$, in the test set. Denote $\hat{\mathbf{M}}$ as the $n \times m$ matrix of estimated GRIn, such that the $j$th column of $\hat{\mathbf{M}}_j$ is $\text{GRIn}_{m_j}$ across all $n$ samples.

Next, we consider the following additive model for the residualized expression of gene $G$:

$$\begin{aligned} \tilde{Y}_G &= \mathbf{M}\beta_{\mathbf{M}} + \varepsilon_{Y_{G_1}}, \\ \tilde{Y}_G - \mathbf{M}\hat{\beta}_{\mathbf{M}} &= \mathbf{X}_G \beta_M + \varepsilon_{Y_{G_2}}. \end{aligned} \tag{2}$$

where $\beta_M$ is the fixed effect-sizes of $\text{GRIn}_{m_j}$ on $\tilde{Y}_G$, $\hat{\mathbf{M}}$ is the matrix of estimated GRIn for all $m_j$ mediators, $\mathbf{X}_G$ are the local-SNPs to gene $G$, and $w_G$ are the "random" or regularized effect sizes of the local-SNPs. We first estimate $\hat{\beta}_M$ by traditional ordinary least squares, where $\hat{\beta}_M = \left(\hat{\mathbf{M}}'\hat{\mathbf{M}}\right)^{-1}\hat{\mathbf{M}}'E_g$. Next, using one of the methods outlined above when estimating $\hat{w}_{M_j}$, we can estimate the effect sizes $\hat{w}_G$ of the local-SNPs on $\tilde{Y}_G$, residualized with $\hat{\mathbf{M}}$, using either elastic net or linear mixed models [3, 4].

#### 1.2 Transcriptomic imputation with MeTWAS

In an external GWAS panel, if individual SNPs are available, we construct the mediator-enriched genetically regulated expression (MeGReX) of gene $G$ directly using $\hat{w}_G$ and $(\hat{w}_j, \hat{\beta}_j),\ 1 \leq j \leq m_G$:

$$MeGReX_G = \sum_{j=1}^{m_G} \mathbf{X}^*_{M_j} \hat{w}_{M_j} \hat{\beta}_{M,j} + \mathbf{X}^*_G \hat{w}_G,$$

where $\mathbf{X}^*_{M_j}$ and $\mathbf{X}^*_G$ are the SNPs in the GWAS panel local to mediator $j$ and gene $G$, respectively. $MeGREX_G$ can be used in downstream tests of association.

### 2 Distal-eQTL prioritization via mediation analysis (DePMA)

#### 2.1 Transcriptomic prediction using DePMA

Expression prediction in DePMA hinges on assessing distal-eSNPs for inclusion in the design matrix via mediation analysis, adopting methods from previous studies [6, 7, 8]. We first split data for gene expression, SNP dosages, and any potential mediators into $k$ training-testing splits. Based on the minor allele frequencies of SNPs and sample size, we recommend a low number of splits (i.e. $k \leq 5$).

In the training set, we identify mediation test triplets that consist of (1) a gene of interest $G$ with expression $Y_G$ (scaled to zero mean and unit variance), (2) a distal-eSNP $s$ in association with $G$ at a user-defined $P$-value threshold (default of $P < 10^{-6}$) with dosages $X_s$, and (3) a set of $m$ biomarkers local to $s$ that are associated with $s$ at a user-defined $P$-value threshold (default of FDR-adjusted $P < 0.05$) with intensities in the $m$ columns of $\mathbf{M}$. The columns of $\mathbf{M}$ are scaled to zero mean and unit variance. Consider the following mediation model:

$$\begin{aligned} Y_G &= X_s\beta_s + \mathbf{M}\beta_{\mathbf{M}} + \mathbf{X}_C\beta_C + \varepsilon_{Y_G} \\ M_j &= X_s\alpha_{M_j} + \mathbf{X}_C\alpha_{C,j} + \varepsilon_{M_j}, \quad 1 \leq j \leq m. \end{aligned} \tag{3}$$

Here, we have $\beta_{\mathbf{M}}$ as the effects of the $M$ mediators local to $s$ on $Y_G$ adjusting for the effects from $s$ and the covariates $\mathbf{X}_C$, and $\alpha_{\mathbf{M}} = (\alpha_{M_1}, \ldots, \alpha_{M_m})'$ as the effects of $s$ on mediators $M_j$, for $1 \leq j \leq m$. We assume that $\varepsilon_{Y_G} \sim N(0, \sigma^2)$ and $\varepsilon_{\mathbf{M}} \sim \mathbf{N}_m(0, \mathbf{\Sigma}_M)$, where $\mathbf{\Sigma}_M$ may have non-zero off-diagonal elements that represent non-zero covariance between mediator intensities. Further, we assume that $\varepsilon_{Y_G}$ and $\varepsilon_{\mathbf{M}}$ are independent. We define the total mediation effect (TME) [9] of SNP $s$ as

$$TME = \alpha^T_{\mathbf{M}}\beta_{\mathbf{M}}.$$

We are interested in SNPs with large absolute TME, which we assess with a two-sided test of $H_0 : TME = 0$. We assess this hypothesis with a permutation test to obtain a permutation $P$-value, as more direct methods of computing standard errors for the estimated TME are often biased [10, 7], We also provide an option to estimate an asymptotic approximation to the standard error of $TME$ and conduct a Wald-type test for $H_0 : TME = 0$. This asymptotic option is significantly faster at the cost of inflated false positives (see **Supplemental Figure S8**). Corresponding to the $t$ testing triplets identified, we obtain vectors of length $t$ of TMEs and $P$-values for each distal-eSNP to $G$. We estimate the $q$-value for each test to adjust for multiple testing [11]. For the predictive model, we select distal SNPs with $TME \neq 0$ at a given $q$-value threshold ($q < 0.10$ as a default) and include them with all local SNPs in a design matrix. We then find estimated SNP weights using either elastic net or weighted least squared regression.

##### 2.1.1 Asymptotic test of total mediation effect

In DePMA, a distal-eQTL $s$ is tested for its total mediation effect on gene $G$ through $m$ mediators that are local to $s$. Consider the following mediation model for $1 \leq j \leq m$:

$$\begin{aligned} Y_G &= X_s \beta_s + \mathbf{M}\beta_{\mathbf{M}} + \mathbf{X}_C \beta_C + \varepsilon_{Y_G} \\ M_j &= X_s \alpha_{M_j} + \mathbf{X}_C \alpha_{C,j} + \varepsilon_{M_j} \end{aligned} \tag{4}$$

.

Here, we construct the total mediation effect

$$\text{TME} = \alpha_{\mathbf{M}}^T \beta_{\mathbf{M}} = \sum_{i=1}^{m} \alpha_{M_i} \beta_{M_i}.$$

Note that TME is distributed as the product of two multivariate Normal distributions. By the multivariate Delta method [12], we can obtain the standard error for the estimated TME. Let $\boldsymbol{\theta} = (\alpha_{\mathbf{M}}, \beta_{\mathbf{M}})$ and define $f(\boldsymbol{\theta}) = TME = \sum_{i=1}^{m} \alpha_{M_i} \beta_{M_i}$.

The first order partial derivative of $f(\hat{\boldsymbol{\theta}})$ is

$$d_{\hat{\boldsymbol{\theta}}} = \frac{\partial(\sum_{i=1}^{m} \alpha_{M_i} \beta_{M_i})}{\partial \hat{\boldsymbol{\theta}}} = [\beta_{\mathbf{M}} \ \alpha_{\mathbf{M}}]^T.$$

We also obtain the estimated variance-covariance matrix $\hat{\boldsymbol{\Sigma}}$ of $\hat{\boldsymbol{\theta}}$:

$$\hat{\mathbf{\Sigma}} = \begin{bmatrix} \hat{\mathbf{\Sigma}}_{\alpha_{\mathbf{M}}} & \hat{\mathbf{\Sigma}}_{\alpha_{\mathbf{M}}\beta_{\mathbf{M}}} \\ \hat{\mathbf{\Sigma}}_{\alpha_{\mathbf{M}}\beta_{\mathbf{M}}} & \hat{\mathbf{\Sigma}}_{\beta_{\mathbf{M}}} \end{bmatrix},$$

where $\hat{\mathbf{\Sigma}}_{\alpha_{\mathbf{M}}}, \hat{\mathbf{\Sigma}}_{\beta_{\mathbf{M}}}$, and $\hat{\mathbf{\Sigma}}_{\alpha_{\mathbf{M}}\beta_{\mathbf{M}}}$ are the variances and covariance of $\hat{\alpha}_{\mathbf{M}}$, $\hat{\beta}_{\mathbf{M}}$, and between $\hat{\alpha}_{\mathbf{M}}$ and $\hat{\beta}_{\mathbf{M}}$, respectively. Sobel previously has shown, that with sufficient sample size, $\hat{\mathbf{\Sigma}}_{\alpha_{\mathbf{M}}\beta_{\mathbf{M}}} \approx 0$ [9, 13]. Thus, the standard error of $\hat{\boldsymbol{\theta}}$ is given by

$$\hat{\sigma}^2_{\hat{\boldsymbol{\theta}}} = d^T_{\hat{\boldsymbol{\theta}}} \hat{\mathbf{\Sigma}} d_{\hat{\boldsymbol{\theta}}}.$$

We can then test $H_0$ : TME $= 0$ against $H_1$ : TME $\neq 0$ with the two-sided Wald-type test with the test statistic $Z = \frac{\alpha^T_{\mathbf{M}}\beta_{\mathbf{M}}}{\sqrt{\hat{\sigma}^2_{\hat{\boldsymbol{\theta}}}}}$ and comparing to the null standard Normal distribution.

We illustrate the trade-off between power and computational speed using the asymptotic Sobel test and the permutation speed. Consider the following simulation framework with $m = 5$ mediators, 3 covariates and a sample size of $n \in \{200, 500, 700, 1000\}$ for the model in Equations 4:

- an $n$-length genotype vector for SNP $s$ is drawn from Binomial$(2, MAF)$, where the minor allele frequency $MAF$ is set at 0.1 in **Supplemental Figure S2**;
- Under the alternative, we simulated $\beta_X \sim N(0,1)$, $\beta_{\mathbf{M}} \sim \mathbf{N}_5(\mathbf{0}, \mathbf{I}_5)$, $\beta_C \sim \mathbf{N}_3(\mathbf{0}, \mathbf{I}_3)$, $\alpha_{M_j}|_{j=1}^{m=5} \sim N(0,1)$, $\alpha_C \sim \mathbf{N}_5(\mathbf{0}, \mathbf{I}_5)$.
- Under the null, all regression parameters were simulated as in the alternative case. However, we set $\alpha_{M_j} = 0|_{j=1}^{m}$ and $\beta_{\mathbf{M}} = \mathbf{0}$.
- Lastly, $\varepsilon_{Y_G} \sim N(0, 1-h^2)$ and $\varepsilon_{M_j} \sim N(0, 1-h^2_M)$, where $h^2 = h^2_M = 0.1$ in **Supplemental Figure S2** below.
- We then constructed $Y_G$ and $\mathbf{M}$ using Equations 4.

We found, that over 10,000 simulations, the permutation test was considerably more powerful, albeit considerably slower. However, in most cases of implementing DePMA, the number of tests of mediation are usually on the order of $10^1$ to $10^2$. We recommend the permutation test in most cases, unless gene $G$ has thousands of identified distal-eQTLs. Parallel implementations have been offered as options in the MOSTWAS package.

#### 2.2 Transcriptomic imputation with DePMA

In an external GWAS panel, if individual SNPs are available, we construct the genetically regulated expression (GReX) of gene $G$ directly using $\hat{w}_G$ and $\hat{w}_t$:

$$GReX_G = \mathbf{X}_t^* \hat{w}_t + \mathbf{X}_G^* \hat{w}_G,$$

where $\mathbf{X}_t^*$ is the matrix of dosages of the $t$ distal-SNPs and $\mathbf{X}_G^*$ is the matrix of dosages of the local SNPs to gene $G$ in the external GWAS panel. $GREX_G$ can be used in downstream tests of association. If individual SNPs are not available, the weighted burden test can be employed using summary statistics with permutation follow-up test [2].

### 3 Tests of associations

If individual SNPs are not available, then the weighted burden $Z$-test proposed by Gusev *et al.* can be employed [2] using summary statistics. Briefly, we compute

$$\tilde{Z} = \frac{\mathbf{W}Z}{(\mathbf{W}\Sigma_{s,s}\mathbf{W}^T)^{1/2}}. \tag{5}$$

Here, $Z$ is the vector of $Z$-scores of SNP-trait associations for SNPs used in estimating $\hat{w}_{M_j}$ and $\hat{w}_G$. The matrix $\mathbf{W}$ is defined as $\Sigma_{e,s}\Sigma_{s,s}^{-1}$, the product of the covariance matrix between all SNPs and the expression of gene $G$ and the covariance matrix among all SNPs. These covariance matrices are estimated from the eQTL reference panel used to estimate $\hat{w}_{M_j}$ and $\hat{w}_G$. The test statistic $\tilde{Z}$ can be compared to the standard Normal distribution for inference. We implement a permutation test conditioning on the GWAS effect sizes to assess whether the same distribution of $\hat{w}_G$ effect sizes could yield a significant association by chance [2]. We permute $\hat{w}_G$ 1,000 times without replacement and recompute the weighted burden test statistic to generate a permutation null distribution for $\tilde{Z}$. This permutation test is only conducted for overall associations at a user-defined significance level (default to FDR-adjusted $P < 0.05$).

Lastly, we also implement a test to assess the information added from distal-eSNPs in the weighted burden test beyond what we find from local SNPs. This test is analogous to a group added-last test in regression analysis, applied here to GWAS summary statistics. Let $Z_l$ and $Z_d$ be the vector of $Z$-scores from GWAS summary statistics from local and distal-SNPs identified by a MOSTWAS model. The local and distal-SNP effects from the MOSTWAS model are represented in $\mathbf{w}_l$ and $\mathbf{w}_d$. Formally, we test whether the weighted $Z$-score $\tilde{Z}_d \equiv \mathbf{w}_d^T Z_d$ from distal-SNPs is significantly larger than 0 given the observed weighted $Z$-score from local SNPs $\tilde{Z}_l \equiv \mathbf{w}_l^T Z_l$, drawing from the assumption that $(\tilde{Z}_l, \tilde{Z}_d)$ follow a bivariate Normal distribution. Namely, we conduct a two-sided Wald-type test for the null hypothesis:

$$H_0 : \mathbf{w}_d^T \mathbf{Z}_d | \mathbf{w}_l^T \mathbf{Z}_l = \tilde{Z}_{l,\text{obs}} = 0.$$

Under the null hypothesis, we can derive that $\tilde{Z}_d | \tilde{Z}_l = \tilde{Z}_{l,\text{obs}}$ is normally distributed with mean and variance determined from the observed local $\tilde{Z}_l$-score, the SNP-effect size vectors $\mathbf{w}_l$ and $\mathbf{w}_d$, and components of the linkage disequilibrium as estimated from the reference panel [14]. Full details and derivation for this added-last test are provided here.

#### 3.1 Added-last test of distal-SNPs

Here, we propose a test to assess the information added from distal-eSNPs in the weighted burden test beyond what we find from local SNPs. Let $\mathbf{Z}_l$ (an $n_l$-vector) and $\mathbf{Z}_d$ (an $n_d$-vector) be the $Z$-scores local and distal SNPs identified by a MOSTWAS model, with $\mathbf{Z} = [\mathbf{Z}_l \ \mathbf{Z}_d]^T$ (an $n$ vector). The local and distal SNP effects from the MOSTWAS model are represented in $\mathbf{w}_l$ (an $n_l$-vector) and $\mathbf{w}_d$ (an $n_d$-vector), with $\mathbf{w} = [\mathbf{w}_l \ \mathbf{w}_d]^T$ (an $n$ vector). Here, we are interested in testing

$$H_0 : \mathbf{w}_d^T \mathbf{Z}_d | \mathbf{w}_l^T \mathbf{Z}_l = \tilde{Z}_{l,\text{obs}} = 0,$$

where $\tilde{Z}_{l,\text{obs}}$ is the observed weighted $Z$-score from local SNPs.

Under the null distribution, as proposed by Pasaniuc et al and Gusev et al in the Imp-G framework[14, 2], we assume that $Z \sim N_n(\mathbf{0}, \mathbf{\Sigma})$, where

$$\mathbf{\Sigma} = \begin{bmatrix} \mathbf{\Sigma}_l & \mathbf{\Sigma}_{l,d} \\ \mathbf{\Sigma}_{l,d}^T & \mathbf{\Sigma}_d \end{bmatrix}$$

is the LD matrix for the SNPs, as estimated from the reference panel. $\mathbf{\Sigma}_l$ and $\mathbf{\Sigma}_d$ represent the LD matrices for local and distal SNPs, respectively. The LD matrix between local and distal SNPs $\mathbf{\Sigma}_{l,d}$ can be assumed to be zero, though recent studies have showed long-range LD in the human genome [15, 16]. We allow the user to set cross-chromosomal LD to 0, though by default, we estimate LD from the reference panel.

Now, we see that, under this null hypothesis, the joint distribution of $(\tilde{Z}_l, \tilde{Z}_d) = (w_l^T Z_l, w_d^T Z_d)$ is given by:

$$\begin{pmatrix} \tilde{Z}_l \\ \tilde{Z}_d \end{pmatrix} \sim N_2 \left( \mathbf{0}, \begin{bmatrix} w_l^T \mathbf{\Sigma}_l w_l & w_l^T \mathbf{\Sigma}_{l,d} w_d \\ w_d^T \mathbf{\Sigma}_{l,d}^T w_l & w_d^T \mathbf{\Sigma}_d w_d \end{bmatrix} \right).$$

It follows that, given $\tilde{Z}_l = \tilde{Z}_{l,\text{obs}}$,

$$\tilde{Z}_d | \tilde{Z}_l = \tilde{Z}_{l,\text{obs}} \sim N \left( \frac{w_l^T \mathbf{\Sigma}_{l,d} w_d}{w_l^T \mathbf{\Sigma}_l w_l} \tilde{Z}_{l,\text{obs}}, w_d^T \mathbf{\Sigma}_d w_d - \frac{[w_l^T \mathbf{\Sigma}_{l,d} w_d]^2}{w_l^T \mathbf{\Sigma}_l w_l} \right).$$

We can use this null distribution for the one-sided test of $H_0 : \mathbf{w}_d^T \mathbf{Z}_d | \mathbf{w}_l^T \mathbf{Z}_l = \tilde{Z}_{l,\text{obs}} = 0$ against $H_1 : \mathbf{w}_d^T \mathbf{Z}_d | \mathbf{w}_l^T \mathbf{Z}_l = \tilde{Z}_{l,\text{obs}} > 0$. These test is implemented in MOSTWAS as a follow-up to the weighted-burden test.

### 4 Simulation framework

We first conducted simulations to assess the predictive capability and power to detect gene-trait associations under various phenotype heritability ($h_p^2$), local heritability of expression ($h_{e,l}^2$), distal heritability of expression ($h_{e,d}^2$), and proportion of causal local ($p_{c,l}$) and distal ($p_{c,e}$) SNPs for MeTWAS and DePMA. We considered two scenarios for each combination of $(h_p^2, h_{e,l}^2, h_{e,d}^2, p_{c,l}, p_{c,e})$: (1) the leveraged association between the distal-SNP and gene of interest exists in both the reference and imputation panel, and (2) the leveraged association between distal-SNP and gene of interest exists in the reference panel but is null in the imputation panel.

Using TCGA data, we extracted 2,592 SNPs local to the gene *ESR1* on Chromosome 6 and 1,431 SNPs local to the gene *FOXA1*. We generated (1) a reference panel with sample size 400 with simulated SNPs, expressions, and one mediator and (2) a GWAS panel of 1,500 samples with simulated SNPs and phenotypes using the following data generating process, modified from Mancuso *et al.*'s framework [17]:

We estimated the linkage disequilibrium $LD$ matrix of the SNPs $X_g$ with $n$ samples and $p$ SNPs, as follows with regularization to ensure $LD$ is positive semi-definite:

$$LD = \frac{1}{n} X_g^T X_g + \frac{1}{10} I_p.$$

We computed the Cholesky decomposition of $LD$ for faster sampling [17]. We simulated SNPs for a 400-sample reference panel $X_{g,ref}$ and 1,500-sample GWAS panel $X_{g,GWAS}$.

We then simulated effect sizes for $p_{c,l}$ of the 2,592 local SNPs $w_{g,l}$ from a standard Normal distribution. We generated locally heritable expression

$$E_{g,l} = X_{g,ref} w_{g,l} + \varepsilon_l,$$

with $\varepsilon_l \sim N(0, 1 - h^2_{e,l})$ and $w_{g,l}$ scaled to ensure the given $h^2_{e,l}$. Similarly, we simulated effect sizes for $p_{c,d}$ of the 1,431 distal-SNPs $w_{g,d}$ and generated the distally heritable intensity of the mediator $M_{g,d}$. We constructed the distally heritable expression $E_{g,d}$ by scaling $M_{g,d}$ by $\beta \sim N(0, 1)$ and adding random noise that scales distal heritability to $h^2_{e,d}$. We lastly formed the total expression $E_g = E_{g,l} + E_{g,d}$.

Next, we simulated the phenotype in the GWAS panel using the GREx as estimated from causal eQTLs to match the variance explained due to genetics. Here, we construct an unobserved heritable expression $E_{g,GWAS}$ for the GWAS panel using the local and distal SNP effect sizes $w_{g,l}, w_{g,d}$, respectively:

$$E_{g,GWAS} = X^*_{g,local} w_{g,l} + X^*_{g,distal} w_{g,d} \beta,$$

where $X^*_{g,\cdot}$ represents genotypes from the GWAS panel. We then drew a causal effect size $\alpha$ for gene expression to simulate a continuous phenotype such that $\alpha \sim N(0, 1)$. We added environmental noise $\varepsilon_* \sim N(0, 1 - h^2_p)$ to scale the total heritability of the phenotype to $h^2_p$.

Here, we also considered a "null" case as well, where the simulated distal eQTLs do not contribute to the simulated phenotype in the GWAS panel (i.e. $w_{g,d} = 0$ for all distal-SNPs). GWAS summary statistics were computed in this step for downstream weighted burden testing. We then fitted predictive models using MeTWAS, DePMA, and local-only models (i.e. FUSION [2]), computed the adjusted predictive $R^2$ in the reference panel, and tested the gene-trait association in the GWAS panel using a weighted burden test. The association study power was defined as the proportion of gene-trait association tests with $P < 2.5 \times 10^{-6}$, the Bonferroni-corrected significance threshold for testing 20,000 independent genes. With these simulated datasets, we also assessed the power of the distal added-last test by computing the proportion of significant distal associations conditional on the local association at FDR-adjusted $P < 0.05$.

[5] Jian Yang, Andrew Bakshi, Zhihong Zhu, Gibran Hemani, Anna A E Vinkhuyzen, Sang Hong Lee, Matthew R Robinson, John R B Perry, Ilja M Nolte, Jana V van Vliet-Ostaptchouk, Harold Snieder, The LifeLines Cohort LifeLines Cohort Study, Tonu Esko, Lili Milani, Reedik Mägi, Andres Metspalu, Anders Hamsten, Patrik K E Magnusson, Nancy L Pedersen, Erik Ingelsson, Nicole Soranzo, Matthew C Keller, Naomi R Wray, Michael E Goddard, and Peter M Visscher. Genetic variance estimation with imputed variants finds negligible missing heritability for human height and body mass index. *Nature genetics*, 47(10):1114–1120, 10 2015.

[6] Brandon L. Pierce, Lin Tong, Lin S. Chen, Ronald Rahaman, Maria Argos, Farzana Jasmine, Shantanu Roy, Rachelle Paul-Brutus, Harm Jan Westra, Lude Franke, Tonu Esko, Rakibuz Zaman, Tariqul Islam, Mahfuzar Rahman, John A. Baron, Muhammad G. Kibriya, and Habibul Ahsan. Mediation Analysis Demonstrates That Trans-eQTLs Are Often Explained by Cis-Mediation: A Genome-Wide Analysis among 1,800 South Asians. *PLoS Genetics*, 10(12), 12 2014.

[7] Nayang Shan, Zuoheng Wang, and Lin Hou. Identification of trans-eQTLs using mediation analysis with multiple mediators. *BMC Bioinformatics*, 20, 3 2019.

[8] Fan Yang, Jiebiao Wang, Brandon L. Pierce, and Lin S. Chen. Identifying cis-mediators for trans-eQTLs across many human tissues using genomic mediation analysis. *Genome Research*, 27(11):1859–1871, 11 2017.

[9] Michael E. Sobel. Asymptotic Confidence Intervals for Indirect Effects in Structural Equation Models. *Sociological Methodology*, 13:290, 1982.

[10] David P. Mackinnon, Chondra M. Lockwood, and Jason Williams. Confidence Limits for the Indirect Effect: Distribution of the Product and Resampling Methods. *Multivariate Behavioral Research*, 39(1):99–128, 2004.

[11] John D. Storey, Andrew J. Bass, Alan Dabney, David Robinson, and Gregory Warnes. qvalue: Q-value estimation for false discovery rate control, 2019.

[12] Yvonne M. Bishop, Stephen E. Fienberg, and Paul W. Holland. *Discrete Multivariate Analysis: Theory and Practice.* Springer, New York, 1975.
