## Supplemental Tables for "MOSTWAS: Multi-Omic Strategies for Transcriptome-Wide Association Studies"

| | **TCGA-BRCA** | **ROS/MAP** |
|---|---|---|
| **Local-only** | 0.037 (0.053) | 0.079 (0.119) |
| **MeTWAS** | 0.040 (0.066) | 0.135 (0.099) |
| **DePMA** | 0.383 (0.194) | 0.405 (0.118) |

Table S1: *Comparison of $h^2$ across local-only, MeTWAS, and DePMA predictive models.* The mean and standard deviation of $h^2$ across all genes that are significantly heritable with the genetic loci considered in the design matrix of each predictive model.

| Gene | Z-statistic (FDR-adjusted $P$) | Cross-validation $R^2$ | TOP GWAS SNP location ($P$) | Permutation FDR-adjusted $P$ | Added last FDR-adjusted $P$ |
|---|---|---|---|---|---|
| **ABCA7** | -1.82 (0.09) | 0.011 | Chr19:553,066 (0.135) | NA | NA |
| **ADAM10** | -1.25 (0.23) | 0.014 | Chr15:59,052,072 ($1.68 \times 10^{-4}$) | NA | NA |
| **APOE** | 2.82 (0.02) | 0.119 | Chr19:45,545,562 ($3.0 \times 10^{-5}$) | $5.0 \times 10^{-3}$ | 0.03 |
| **BIN1** | 1.91 (0.08) | 0.010 | Chr22:24,199,787 ($8.53 \times 10^{-4}$) | NA | NA |
| **CD2AP** | 1.52 (0.15) | 0.011 | Chr6:47,432,637 ($1.23 \times 10^{-4}$) | NA | NA |
| **CLU** | -2.41 (0.04) | 0.012 | Chr8:27,465,312 ($1.33 \times 10^{-4}$) | 0.83 | 0.44 |
| **FERMT2** | 2.13 (0.06) | 0.017 | Chr14:53,305,626 ($1.38 \times 10^{-4}$) | NA | NA |
| **MEF2C** | 2.20 (0.06) | 0.016 | Chr5:88,359,039 (0.020) | NA | NA |
| **PLCG2** | -2.48 (0.04) | 0.010 | Chr16:81,879,218 (0.037) | 0.66 | 0.07 |
| **SORL1** | 2.91 (0.02) | 0.043 | Chr11:121,446,813 (0.032) | 0.04 | $4.5 \times 10^{-3}$ |
| **ZCWPW1** | -4.56 ($6.1 \times 10^{-5}$) | 0.018 | Chr7:100,435,157 (0.074) | 0.03 | $1.3 \times 10^{-5}$ |

Table S2: *Summary statistics for known Alzheimer's risk-associated loci identified by MOSTWAS models.* TWAS associations (weighted $Z$-score and FDR-adjusted $P$-value) with late-onset Alzheimer's risk from GWAS statistics from IGAP. The top IGAP GWAS SNP in the identified loci with its location and $P$-value are provided. For the 6 loci with significant TWAS associations, the FDR-adjusted $P$-value for the follow-up distal SNP added last test is provided.

| Gene | Cross-validation $R^2$ | PGC $Z$-statistic (UKBB $Z$) | Top GWAS SNP location ($P$) | Permutation FDR-adjusted $P$ |
|---|---|---|---|---|
| **ADAD2** | 0.050 | 5.89 (4.16) | Chr5:35,639,107 ($4.05 \times 10^{-3}$) | $3.5 \times 10^{-5}$ |
| **CACNA2D3** | 0.033 | 3.41 (2.88) | Chr7:12,268,243 ($1.27 \times 10^{-2}$) | 0.046 |
| **FAM43B** | 0.035 | -4.03 (-2.85) | Chr2:73,148,399 ($2.09 \times 10^{-2}$) | 0.028 |
| **MGC29506** | 0.022 | 3.51 (5.54) | Chr5:139,536,922 ($1.48 \times 10^{-3}$) | $3.5 \times 10^{-5}$ |
| **OR8U1** | 0.022 | -3.19 (-4.21) | Chr11:56,676,947 ($4.90 \times 10^{-5}$) | 0.049 |
| **SYT1** | 0.015 | -5.58 (-3.16) | Chr7:12,269,417 ($1.29 \times 10^{-2}$) | 0.040 |
| **YJEFN3** | 0.010 | 5.82 (7.22) | Chr7:12,276,011 ($1.35 \times 10^{-2}$) | 0.038 |

Table S3: *Summary statistics for 7 MDD risk-associated loci identified by MOSTWAS models.* TWAS associations with major depressive disorder from GWAS statistics from Psychiatric Genomics Consortium that were replicated with GWAX summary statistics in UK Biobank. The top PGC GWAS SNP in the identified loci with its location and $P$-value are provided.

| Gene | Cross-validation $R^2$ | iCOGs $Z$-statistic (Added-last $Z$) | Top GWAS SNP location ($P$) | Permutation FDR-adjusted $P$ |
|---|---|---|---|---|
| **C16orf13** | 0.019 | 4.51 (5.18) | Chr3:10720351 (0.13) | 0.03 |
| **C9orf169** | 0.011 | 4.18 (3.97) | Chr19:44949849 (0.04) | 0.03 |
| **CTRL** | 0.051 | 4.51 (3.88) | Chr10:2798136 (0.06) | 0.03 |
| **DNAL4** | 0.013 | -3.94 (-4.57) | Chr22:38681840 (0.01) | 0.04 |
| **LOC221710** | 0.014 | 5.34 (5.08) | Chr1:152983865 ($8.5 \times 10^{-4}$) | 0.03 |
| **MAP3K6** | 0.021 | -4.10 (-4.00) | Chr1:27686314 (0.01) | 0.05 |
| **MAP4K5** | 0.020 | 3.76 (1.26) | Chr14:50502944 ($1.3 \times 10^{-4}$) | 0.03 |
| **NPAT** | 0.115 | -3.92 (-3.72) | Chr20:4217738 (0.02) | 0.04 |
| **RPLP1** | 0.040 | -3.82 (-3.83) | Chr18:1592917 (0.18) | $1.4 \times 10^{-4}$ |
| **SPATA5L1** | 0.042 | 3.76 (No distal SNPs in model) | Chr15:45593323 (0.01) | $1.4 \times 10^{-4}$ |
| **TXNRD2** | 0.047 | 3.91 (4.64) | Chr22:19735425 ($3.5 \times 10^{-3}$) | 0.05 |

Table S4: *Summary statistics for 11 breast cancer-specific survival-associated loci identified by MOSTWAS models.* TWAS associations with breast cancer survival from GWAS statistics from iCOGs with permutation test results and added-last $Z$-statistics. The top iCOGs GWAS SNP in the identified loci with its location and $P$-value are provided.
